# Amphipathic Bax core dimer forms part of apoptotic pore wall in the mitochondrial membrane

**DOI:** 10.1101/2020.12.18.423546

**Authors:** Fujiao Lv, Fei Qi, Zhi Zhang, Maorong Wen, Alessandro Piai, Lingyu Du, LiuJuan Zhou, Yaqing Yang, Bin Wu, Zhijun Liu, Juan del Rosario, James J. Chou, David W. Andrews, Jialing Lin, Bo OuYang

**Affiliations:** State Key Laboratory of Molecular Biology, Shanghai Institute of Biochemistry and Cell Biology, CAS Center for Excellence in Molecular Cell Science, Chinese Academy of Sciences, Shanghai 201203, China; University of Chinese Academy of Sciences, Beijing 100049, China; Department of Biochemistry and Molecular Biology, University of Oklahoma Health Sciences Center, 940 Stanton L. Young Boulevard, Oklahoma City, OK 73126, USA; Department of Biological Chemistry & Molecular Pharmacology, Harvard Medical School, Boston, MA 02115, USA; National Facility for Protein Science in Shanghai, Chinese Academy of Sciences, Shanghai 201210, China; Biological Sciences, Sunnybrook Research Institute, Toronto, Ontario, Canada, M4N 3M5; Stephenson Cancer Center, 800 Northeast 10th Street, Oklahoma City, OK 73104, USA

**Keywords:** Bax core dimer, NMR, pore formation, lipid bilayer, functional mutagenesis

## Abstract

Bax proteins form pores in the mitochondrial outer membrane to initiate apoptosis. They may embed in the cytosolic leaflet of the lipid bilayer generating tension to induce a lipid pore with radially arranged lipids forming the wall. Alternatively, they may comprise part of the pore wall. However, there is no unambiguous structural evidence for either hypothesis. Using NMR, we determine a high-resolution structure of the Bax core region that forms a dimer with the nonpolar surface covering the lipid bilayer edge and the polar surface exposed to water. The dimer tilts from the bilayer normal, not only maximizing nonpolar interactions with lipid tails but creating polar interactions between charged residues and lipid heads. Structure-guided mutations demonstrate the importance of both protein-lipid interactions in Bax pore assembly and core dimer configuration. Therefore, the Bax core dimer forms part of the proteolipid pore wall to permeabilize mitochondria.

## Introduction

Apoptotic cell death initiates when the mitochondrial outer membrane (MOM) is permeabilized by Bax or Bak, two pore-forming proteins in the Bcl-2 family. Bax and Bak share the same structural fold with the colicine family of α-helical pore-forming bacterial toxins (Moldoveanu et al., 2006; Parker et al., 1989; Suzuki et al., 2000). Despite more than three decades of intensive investigation, it is still unclear how these proteins form pores in various membranes and what are the pore structures that allow molecules of various sizes to cross diverse membranes (Andrews, 2014; Cosentino and Garcia-Saez, 2017; Dal Peraro and van der Goot, 2016; Uren et al., 2017). In particular, it is unclear how Bax can form giant pores in the MOM that can be tens to hundreds of nanometers in diameter (Grosse et al., 2016; Salvador-Gallego et al., 2016). The most popular proteolipid pore model requires Bax to fulfill two functions (Cosentino and Garcia-Saez, 2017). First, Bax would embed in the cytosolic leaflet of the MOM lipid bilayer, asymmetrically expanding it to bend the bilayer and generate membrane tension that eventually breaks the lipid bilayer. The lesion in the membrane exposes the nonpolar lipid acyl tails to aqueous milieu, and hence, is thermodynamically unstable. An immediate solution is to cover the exposed acyl tails of the bilayer lipids with non-bilayer lipids, for example, lipids that are radially arranged to a micellar structure, resulting in an aqueous pore lined by the polar or charged heads of bent lipids. However, bending the lipids at the pore rim generates line tension that is proportional to the pore radius, and hence, opposes pore opening. Thus, the second function of Bax is to reduce the line tension by intercalating between the bent lipids and reinforcing the pore rim around the bilayer lipids.

In healthy cells Bax is mostly a soluble cytoplasmic protein but occasionally visiting the mitochondria as a peripherally bound protein (Edlich et al., 2011). To become a membrane protein and fulfill the pore-inducing and stabilizing functions in stressed cells, Bax must change conformation. The conformation change is triggered by BH3 proteins of the Bcl-2 family (Fig. 1A). In particular, transient binding by Bid or Bim unfolds the soluble and globular Bax structure of a bundle of nine α-helices (α1 to α9) (Fig. 1A, soluble Bax) (Suzuki et al., 2000), resulting in an exposed α9 at the C-terminal that can insert into the MOM, and an exposed α2 to α5 region that can interact with its counterpart in another activated and unfolded Bax protein (Chi et al., 2020; Gavathiotis et al., 2010; Gavathiotis et al., 2008; Lovell et al., 2008; Suzuki et al., 2000). According to a crystal structure, the two α2 to α5 regions can form a symmetric and amphipathic dimer with a polar and charged α2-α3 surface on the top and a nonpolar α4-α5 surface at the bottom (Fig. 1A, Bax (α2-α5) dimer) (Czabotar et al., 2013). This structure would fulfill the two functions if the sidechains of the nonpolar α4-α5 residues could not only insert into the cytosolic leaflet of the MOM lipid bilayer to induce the pore opening, as the “in-plane” insertion model postulated (Fig. 1A, in-plane model) (Westphal et al., 2014), but also intercalate between the bent lipids at the pore rim to stabilize the opening, as the “clamp” model suggested (Fig. 1A, clamp model) (Bleicken et al., 2014).

**Figure 1.**
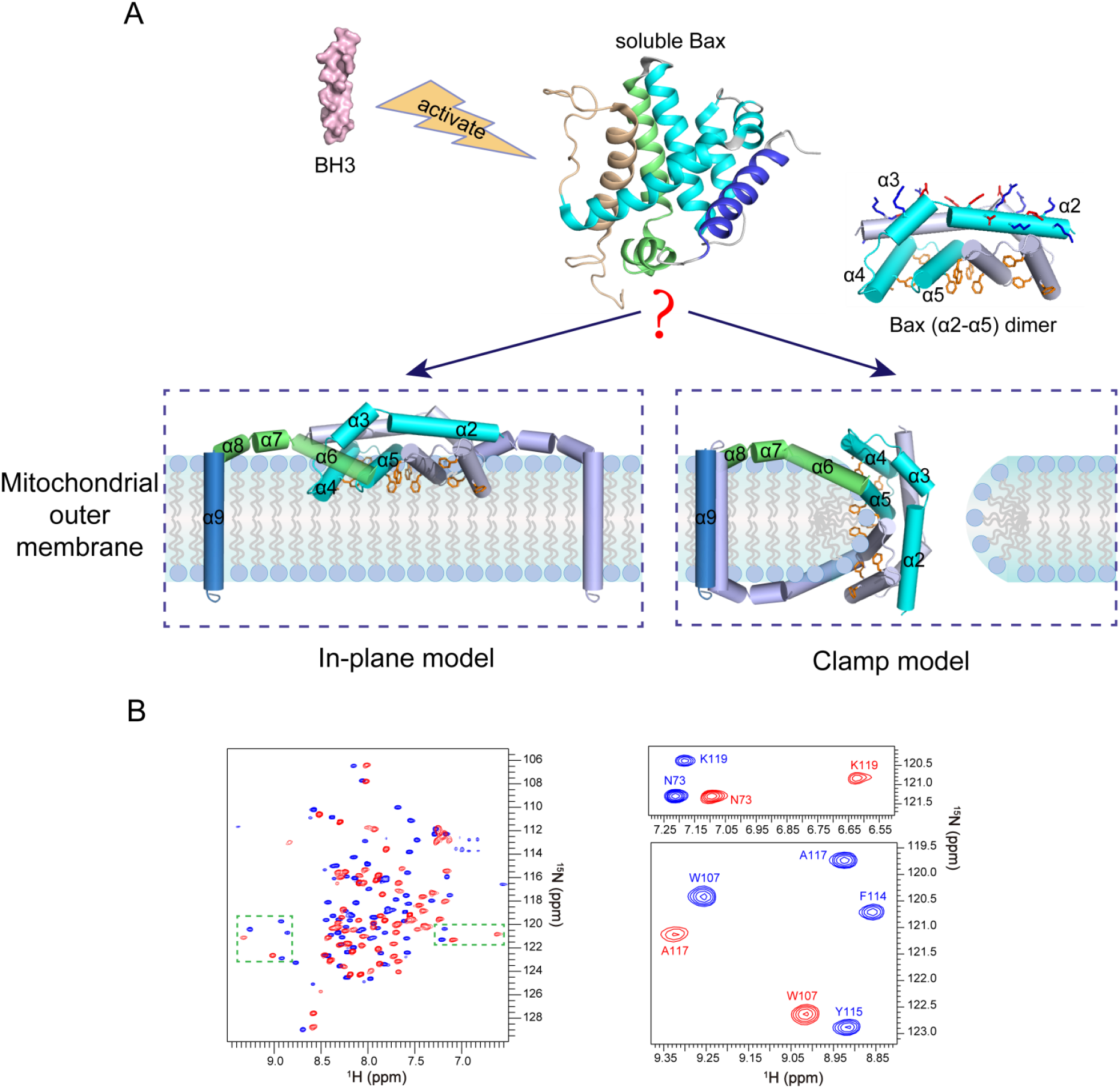
Models for the membrane topology of Bax and NMR spectra of Bax (α2-α5) (A) The “in-plane” and the “clamp” models for the membrane topology of pore-forming Bax protein. After activation by BH3 proteins, the soluble Bax protein unfolds the α-helical bundle structure and inserts the α9 helix into the MOM. The tail-anchored Bax proteins dimerize via their α2, α3, α4 and α5 helices resulting in an amphipathic structure (shown as Bax (α2-α5) dimer with the protomers in cyan or gray color) with a polar surface containing positive and negative charged α2 and α3 residues (blue and red sticks) and a nonpolar surface containing aromatic α4 and α5 residues (orange rings). According to the in-plane model, the α2-α5 dimer engages the planer cytosolic surface of the MOM with the aromatic α4 and α5 residues projected into the cytosolic leaflet of the lipid bilayer to generate membrane tension thereby inducing a lipid pore with radially arranged lipids at the rim. In contrast, the clamp model positions the α2-α5 dimer at the lipid pore rim with the aromatic α4 and α5 residues intercalated between the radiated lipids to reduce line tension thereby stabilizing the pore. As illustrated, the membrane topologies of other helices are also different between the models. The structures of the soluble Bax protein and the α2-α5 dimer were generated from PDB files 1F16 and 4BDU, respectively, using PyMOL program. The structures of other helices are modeled based on the corresponding structures in the soluble Bax protein. (B) 2D ^1^H-^15^N TROSY-HSQC spectra of DMPC/DHPC bicelle-bound Bax (α2-α5) (blue) and soluble Bax (α2-α5) (red) at 600 MHz. The magnified regions highlight the spectral changes for some residues in the bicelle-bound and soluble proteins.

Accumulating biochemical and biophysical evidence has supported these structure-based functions. (i) Site-specific crosslinking, double electron-electron resonance (DEER) spectroscopy and Förster resonance energy transfer have verified the dimeric interaction between the α2 to α5 regions of two full-length Bax proteins integrated into the native or model mitochondrial membrane (Bleicken et al., 2014; Dewson et al., 2012; Gahl et al., 2014; Zhang et al., 2016). (ii) Accessibility of cysteine residues placed in the nonpolar surface of the α2 to α5 dimer to a charged sulfhydryl reactive labeling agent (IASD), electron paramagnetic resonance (EPR) detected accessibility of spin probes attached to the nonpolar surface to molecular oxygen enriched in the lipid bilayer or paramagnetic quenchers soluble in water, fluorescence emission and quenching of environment sensitive dyes attached to the nonpolar surface, and H-D exchange of the α4 to α5 region in a membrane-bound Bax oligomer are consistent with their locations within the membrane (Annis et al., 2005; Bleicken et al., 2018; Bleicken et al., 2010; Flores-Romero et al., 2017; Hauseman et al., 2020; Westphal et al., 2014; Zhang et al., 2016). (iii) In contrast to the stable conformation of the α2 to α5 dimer, the DEER spectral data from the α6 to α9 region indicate a dynamic conformation that would allow the α6 from one protomer to engage the cytosolic surface of the MOM, and the α6 from the other protomer to engage the intermembrane surface, with the two α9 membrane anchors in opposite orientations (Bleicken et al., 2014). This topographic model provides a mechanism that would clamp the α2 to α5 dimer at the pore rim using the α6 to α9 regions, and fits the concave nonpolar surface of the dimer nicely to the convex micellar lipid surface (Fig 1A, clamp model) (Bleicken et al., 2014).

Therefore, both of the proposed functions for Bax have been supported by experimental evidence. As illustrated in Figure 1A, the pore-inducing function would be fulfilled by the nonpolar surface of the α2 to α5 dimer if it engages the cytosolic leaflet of the MOM lipid bilayer. In contrast, if the α2 to α5 dimer intercalates between the bent lipids surround the opening of a lipid pore, it would stabilize the pore. However, the structure of the α2 to α5 dimer was determined by crystallography in the absence of membranes. Thus, we do not know where and how the dimer engages the membrane. On the other hand, although the IASD-labeling, EPR, fluorescence and H-D exchange data were obtained with fulllength Bax proteins bound to membranes, they do not possess enough structural resolution to differentiate the two potential membrane topologies of the α2 to α5 regions proposed by the in-plane and the clamp models (Andrews, 2014; Bleicken et al., 2018). To overcome these drawbacks, we first determined the structure of the α2 to α5 region bound to a lipid bicelle using NMR. The resulting atomic resolution model for how the α2 to α5 dimer interacts with a model lipid bilayer is more consistent with the clamp model than the in-plane model. We then performed structure-guided mutagenesis to assess how distinct interactions between specific residues within and around the nonpolar surface of the α2 to α5 dimer and lipids in the bilayer contribute to the pore-forming function of intact Bax protein in a model mitochondrial membrane, and to the dimer configuration in the native mitochondrial membrane. Our study demonstrates that the α2 to α5 dimer forms part of the wall that separates the nonpolar lipid bilayer from the aqueous pore via ionic or polar interactions with the negatively charged or polar lipid headgroups and hydrophobic interactions with the nonpolar lipid acyl chains. Thereby, we propose a model for the mitochondrial Bax pore, which is at least partially lined by a series of α2 to α5 dimers that are stitched together by other Bax regions.

## Results

### Structure determination of Bax (α2-α5) in lipid bicelles

To study the structural properties of the Bax core domain in membrane, we determined the structure of the bicelle-bound Bax α2 to α5 region (α2-α5) following a NMR strategy taken earlier for oligomeric membrane proteins (Fu et al., 2019). We made bicelles with 1,2-dimyristoyl-sn-glycero-3-phosphocholine (DMPC) and 1,2-dihexanoyl-sn-glycero-3-phosphocholine (DHPC) at a molar ratio (q) ≥ 0.5 to yield lipid discs with a radius of ~25 Å for the bilayer region (Fig. S1A), large enough to be certain that the Bax protein can be embedded in a lipid bilayer environment. These lipid discs also have radially arranged DHPC lipids at their rim (Chen et al., 2018), similar to the lipid pore rim to which the Bax protein may bind. We expressed and purified the isotope-labeled human Bax (α2-α5) protein (residues D53 to K128 with C62S and C126S mutations). The resulting soluble Bax (α2-α5) is stable enough to generate a two-dimensional (2D) ^1^H-^15^N correlation spectrum with good resolution and dispersion (Fig. 1B). Addition of the lipid bicelles dramatically changed the NMR spectrum of the Bax (α2-α5) (Fig. 1B), indicating that the protein undergoes large conformational changes due to the interaction with membranes. SEC-MALS analysis showed that the soluble Bax (α2-α5) is mainly a tetramer (Fig. S1B), which was detected by cross-linking alongside the dimer and trimer (Fig. S1C). The cross-linking efficiency was low for the bicelle-bound Bax (α2-α5) (Fig. S1C), possibly because the BS^3^ crosslinker is membrane impermeable. Nonetheless, a small fraction of the bicelle-bound protein was crosslinked as a dimer.

To determine the structure, nearly all (97%) of the backbone resonances in the bicelle-bound Bax (α2-α5) were assigned using standard TROSY-based triple resonance experiments (Fig. S2A). Most resonances are clearly shifted from the assigned backbone resonances of the soluble protein (Fig. 1B and S2B). The secondary structures of the bicelle-bound Bax (α2-α5) were derived from analysis of the backbone chemical shifts using the TALOS+ program (Shen et al., 2009). Overall the helical regions of Bax (α2-α5) in bicelles correlate well with those in the crystal structure (PDB code: 4BDU) (Czabotar et al., 2013). We then determined the local structures of the bicelle-bound Bax (α2-α5) monomers and assembled the dimer structures by using 702 local and 60 long-range distance restraints derived from nuclear Overhauser enhancement (NOE) measurements (Fig. S3A-B; Table 1). The 15 lowest-energy structures (PDB code: 6L8V) out of 200 calculated dimer structures converged to a root mean squared deviation (RMSD) of 1.058 Å and 1.504 Å for backbone and all heavy atoms, respectively (Fig. 2A; Table 1). Furthermore, a sulfhydryl reactive nitroxide spin label, MTSL (Fu et al., 2019), was introduced into two single-Cys Bax (α2-α5) mutants (A82C and S126C) to collect the paramagnetic resonance enhancement (PRE) data that provided the inter-monomer distance restraints confirming the structures of the symmetric antiparallel Bax (α2-α5) dimers in the bicelles (Fig. S3C).

**Table 1.** NMR and Refinement Statistics for bicelle-bound Bax (α2-α5) Structures

|  |  |
| --- | --- |
| **NMR distance and dihedral constraints** |  |
| Distance constraints from NOE | 762 |
| Intra-chain NOEs | 702 |
| Inter-chain NOEs | 60 |
| Total dihedral angle restraints | 260 |
| $\Phi$ (TALOS) | 130 |
| $\Psi$(TALOS) | 130 |
| **Structure statistics <sup>a</sup>** |  |
| Violations (mean ± SD) | |
| Distance constraints ($\text{\AA}$) | $0.077 \pm 0.005$ |
| Dihedral angle constraints ($^\circ$) | $0.653 \pm 0.014$ |
| Deviations from idealized geometry |  |
| Bond lengths ($\text{\AA}$) | $0.006 \pm 0.000$ |
| Bond angles ($^\circ$) | $0.310 \pm 0.057$ |
| Improper ($^\circ$) | $0.419 \pm 0.018$ |
| Average pairwise RMSD ($\text{\AA}$) <sup>b</sup> | |
| Heavy atoms | 1.504 |
| Backbone | 1.058 |
| PDB ID | 6L8V |

<sup>a</sup> Statistics are calculated and averaged over an ensemble of the 15 lowest energy structures of the 200 calculated structures.
<sup>b</sup> The precision of the atomic coordinates is defined as the average RMSD between the 15 lowest energy structures and their mean coordinates.

**Figure 2.**
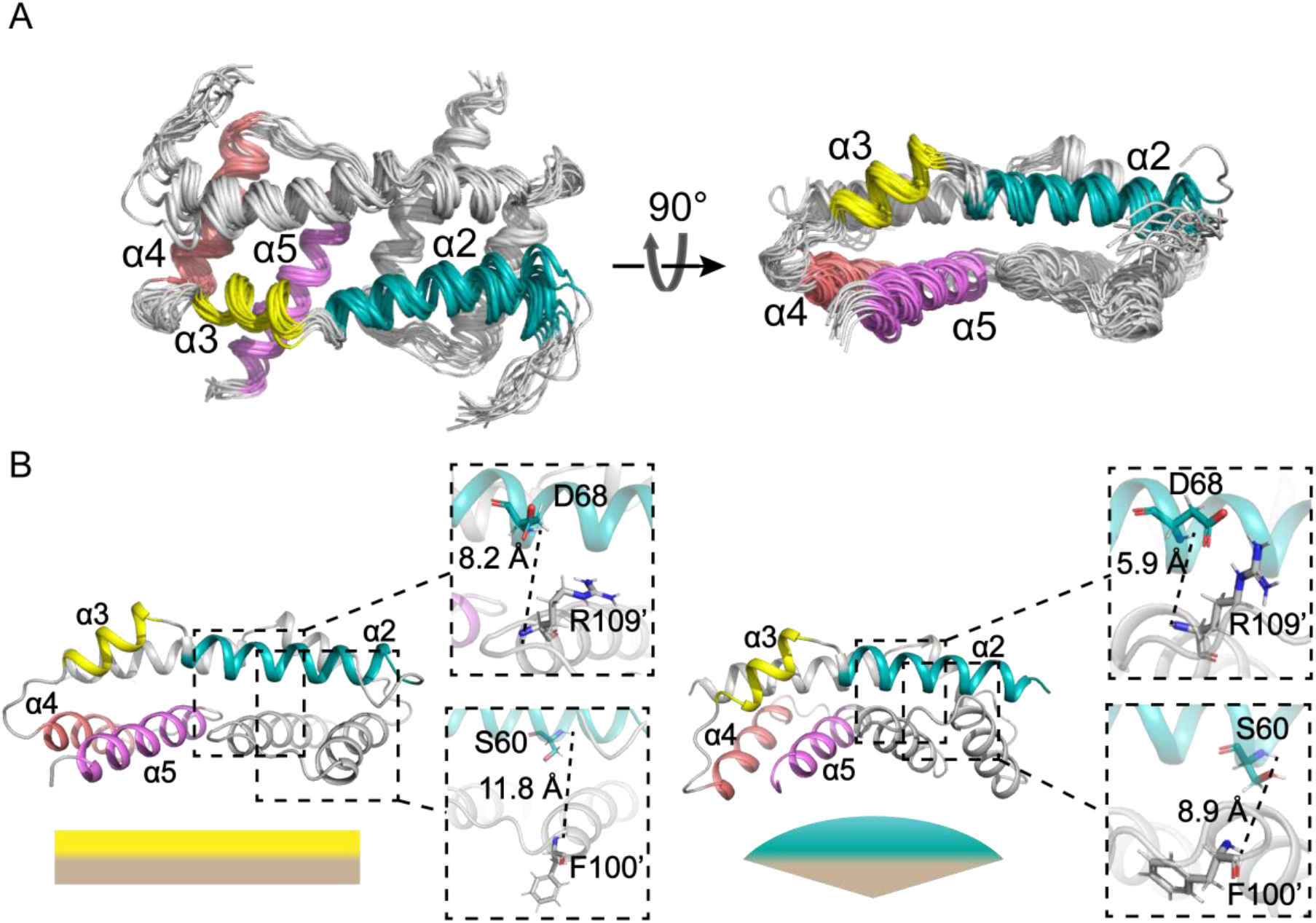
Structures of Bax (α2-α5) in lipid bicelles determined by NMR. (A) Ensemble of 15 lowest-energy structures for Bax (α2-α5) dimer in bicelles calculated from the geometry constraints obtained from the NMR experiments summarized in Table 1. The backbone structures are shown as thin ribbons. (B) Comparison of the NMR structure of Bax (α2-α5) in bicelles (left; PDB code: 6L8V) and the crystal structure (right; PDB code: 4BDU). The NMR or crystal structure has a flat or concave bottom surface formed by the α4-α5 regions, respectively, as indicated by the complimentary flat or convex shapes underneath. The zoom-in regions display the distances between the amide protons of D68-R109’ and S60-F100’, which are longer in the NMR structure than the crystal structure.

The bicelle-bound Bax (α2-α5) dimer structure is different from the previously reported crystal structure (PDB code: 4BDU) (Fig. S4) (Czabotar et al., 2013), as expected from the dramatically different NMR spectra for the bicelle-bound and the soluble Bax (α2-α5) (Fig. 1B). Alignment of the two structures shows a 3.878-Å backbone RMSD (Fig. S4). In both structures, the two α2 helices are antiparallel and each interacts with the α3, α,4 and α5 helices of the other protomer (Fig. 2B). However, upon the bicelle binding, the C-terminus of the α4 helix and the N-terminus of the α5 helix tilt away from the α2 helix to yield a larger gap in the dimerization interface with less inter-molecular interactions (Fig. 2B). As a result, the distances between the amide protons of S60-F100’ and D68-R109’ are increased in the bicelle-bound structure (Fig. 2B). Furthermore, the α4 and α5 helices in one protomer form a hydrophobic surface with their counterparts in the other protomer. The angles between the two α4 helices and between the two α5 helices in the dimeric Bax (α2-α5) are reduced from ~82° and ~26° in the crystal structure to ~12° and ~11° in the bicelle-bound structure, respectively, thereby changing the concave surface to a flat surface (Fig. 2B). Finally, the α5 helix is elongated in the bicelle-bound structure with residues K123 to S126 forming an additional α helical turn, while the structural information of these residues is missing in the crystal structure (Fig. S4).

### Interaction of Bax (α2-α5) with lipid bicelles

To determine how the Bax (α2-α5) dimer interacts with a membrane, we performed protein-lipid NOE experiments to examine the interaction between the dimer and the lipid bicelle. We recorded an ^15^N-edited NOE spectroscopy (NOESY) spectrum using the [^15^N, ^2^H]-labeled Bax (α2-α5) protein reconstituted in the bicelles composed of regular DMPC and deuterated DHPC. The crosspeaks in the NOESY spectrum appear when two protons are close to each other, typically within 5 Å (Skinner and Laurence, 2008). The NOE strip plot taken from the 3D ^15^N-edited NOESY-TROSY-HSQC spectrum shows that the residues in α2 and α3 helices have only water crosspeaks from either NOE or proton exchange or both and no NOEs to the DMPC lipids, indicating that these helices are mainly exposed to water and not interacting with the lipid bilayer (Fig. 3A). Residues T85 and D86 show NOEs only to the headgroups of DMPC lipids, whereas R89 and E90 in α4 helix show NOEs to both headgroups and acyl chains of DMPC lipids, indicating their shallow immersion in the lipid bilayer. In contrast, residues L113, F114, F116 and A117 in α5 helix display strong NOEs to the entire acyl chains of DMPC lipids including the methyl groups at the ends, indicating deep immersion in the lipid bilayer.

**Figure 3.**
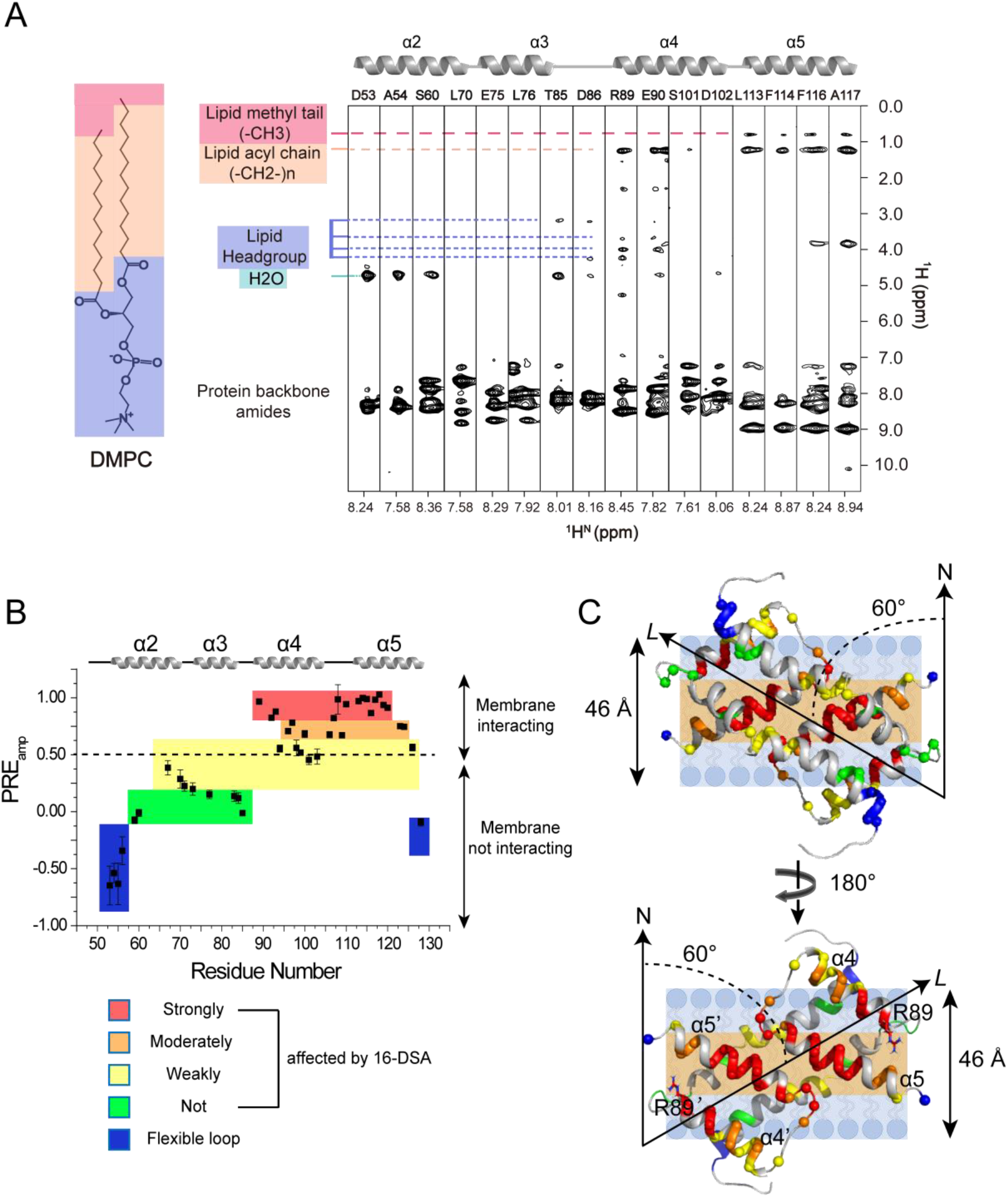
Interaction of Bax (α2-α5) with lipid bicelles. (A) NOE strips taken from 3D ^15^N-edited NOESY-TROSY-HSQC spectrum (200 ms NOE mixing time) recorded at 900 MHz using the [^15^N, ^2^H]-labeled Bax (α2-α5) in DMPC/DHPC bicelles with q = 0.55. The indicated crosspeaks in the aliphatic regions are the NOEs between the protein backbone amide protons and the lipid aliphatic protons. (B) Residue-specific PRE_amp_ of Bax (α2-α5) in DMPC/DHPC bicelles with q = 0.6 determined from the lipophilic PRE analysis. The PRE_amp_ values were derived from the 16-DSA titration. The plot is colored according to the PRE_amp_ values that are proportional to the effects of 16-DSA on the individualresidues of Bax (α2-α5). The residues with the least PRE_amp_ values in the blue colored areas are in the flexible terminal regions. (C) Position of a Bax (α2-α5) dimer structure in ribbon representation relative to the lipid bilayer of a bicelle. The protein structure is placed against the lipid bilayer with a 60° tilt angle between its longest axis (L) and the bilayer normal axis (N) as derived from the best sigmoidal fitting. The hydrophobic α4-α5 surface contacts the lipid bilayer. Amide protons are shown as spheres color-coded similarly as in panel B. The polar headgroup region and the nonpolar acyl tail region of the lipid bilayer are indicated by light blue and light orange colored strips, respectively. The estimated thickness for the DMPC lipid bilayer is indicated on the left side. The Bax (α2-α5) dimer structure on the top is rotated 180° to generate the structure on the bottom showing the nonpolar residues in the α4-α5 surface, most of which contact the nonpolar core of the lipid bilayer. Note that the two positively charged R89 residues on the boundary of the nonpolar surface are close to the polar lipid headgroups (Bottom).

To analyze the depth of immersion of the Bax (α2-α5) dimer along the bicelle normal axis, we used the paramagnetic probe titration (PPT) method that has been applied to study the transmembrane domains of immune receptors and viral proteins (Chen et al., 2018; Piai et al., 2017). Following a published protocol (Fu et al., 2019), we first titrated the water-soluble paramagnetic probe gadolinium (III) 1,4,7,10-tetraazacyclododecane-1,4,7,10-tetraacetate (Gd-DOTA) into the bicelle-bound Bax (α2-α5) to measure residue-specific PRE amplitudes (PRE_amp_). The PRE_amp_ values derived from the analysis of the peak intensity decay versus the paramagnetic probe concentration report residue-specific membrane immersion depths. Unexpectedly, a plot of PRE_amp_ versus residue number shows close PRE_amp_ values ~0.9-1.0 for all the residues, which is not a typical plot expected for transmembrane proteins (Fig. S5A). However, the residues in the α4 and α5 helices display slower intensity decay rate as the concentration of Gd-DOTA increases than the residues in the α2 and α3 helices (examples shown in Fig. S5B), indicating that the α4 and α5 helices are less exposed to the aqueous solvent.

To clearly define the position of Bax (α2-α5) relative to the bilayer, we acquired a set of lipophilic PRE data. The bicelle-bound Bax (α2-α5) was titrated with the membrane-embedded paramagnetic agent 16-doxyl-stearic acid (16-DSA) that places the paramagnetic probe in the center of the lipid bilayer. The residue-specific PRE_amp_ was derived using the same approach as that employed for the hydrophilic PRE. Instead of observing a flat hydrophilic PRE_amp_ plot, the lipophilic PRE_amp_ plot shows an overall bell-shaped profile (Fig. 3B). The N- and C-termini of Bax (α2-α5) exhibit negative PRE_amp_ values, indicating that they are very flexible. Most of α2 and α3 helices exhibit very small PRE_amp_ values (< 0.25), confirming that they are not in direct contact with the bicelle. In contrast, the PRE_amp_ values for part of α4 helix and entire α5 helix are above 0.5 (Fig. 3B), indicating that the residues in these helices are close to the lipid acyl tails, likely interacting with the lipid bilayer directly.

Taken together, the lipophilic PRE and lipid NOE data demonstrate that the α4 and α5 helices in Bax (α2-α5) dimer are directly associated to the DMPC lipid bilayer. However, the flat PRE_amp_ (residue number) plot from hydrophilic PRE indicates that no protein region is in a conventional transmembrane topology which could shield the effect of the hydrophilic paramagnetic agent. To fit all sets of data, we propose a model in which the hydrophobic α4-α5 surface of the Bax (α2-α5) dimer binds to the edge of the lipid bilayer with the α2-α3 hydrophilic surface facing the aqueous solvent (Fig. 3C). Given the unconventional membrane-protein interaction, the PRE effect of Gd-DOTA in the aqueous phase is so strong that affects all the residues to the maximum extent independently of their positions. In contrast, the PRE effect of the lipophilic 16-DSA in the lipid bilayer is more localized and generating a gradient, thus being used for further analysis. Since lipophilic PRE_amp_ is proportional only to the residue position along the bicelle normal, the relative position of each residue to the bilayer center could be estimated by fitting a structural model to the experimental PRE_amp_ values as we described before (Chen et al., 2018; Piai et al., 2017). Thus, we calculated the distance r_z_ for each residue in the Bax (α2-α5) dimer structure to an arbitrary point on the bilayer normal axis (examples shown in Fig. S5C). Then we repeated the sigmoidal fitting of PRE_amp_ (r_z_) data while rotating the protein structure until finding the best-fit model. The R^2^_adj_ versus tilt angle plot (Fig. S5D) indicates that the best-fit model has an angle of ~60° between the protein axis (L) and the bilayer normal (N) (Fig. S5E), suggesting that the protein is docked to the membrane in an orientation illustrated in Figure 3C. This orientation also agrees with the protein hydrophobicity map, as it places most of the hydrophobic residues on the α4-α5 surface in contact with the hydrophobic core of the lipid bilayer.

### Structure validation by functional mutagenesis

The NMR structure of Bax (α2-α5) dimer bound to lipid bicelle provides a high-resolution model for how this Bax core dimer interacts with the lipid bilayer. In particular, many nonpolar residues throughout the α4 and α5 helices and a polar residue, S118, at the middle of α5 helix interact with the nonpolar lipid acyl chains, whereas a positively charged residue, R89, at the N-terminus of α4 helix interacts with the negatively charged lipid head groups (Fig. 4A). These protein-lipid interactions would allow the core dimer to form part of the wall that separates the nonpolar lipid bilayer from the aqueous pore. Mutations of these residues to negatively charged residues are expected to reduce these protein-lipid interactions, thereby impeding the pore-forming function of Bax protein. To test this structurebased functional model for the Bax (α2-α5) dimer, we mutated a representative residue of each category to a negatively charged residue, and determined the interaction of the resulting mutants, R89E, A117D, and S118D, with the lipid bilayer (Fig. 4A). While we were unable to purify Bax (α2-α5) with A117D, we successfully purified Bax (α2-α5) with R89E or S118D and reconstituted each into the bicelles (Fig. S6). Most of the resonances from the α4-α5 residues in both soluble and bicelle-bound S118D mutant were missing in the NMR spectra, preventing us from determining how the mutant interacts with the membrane since these helices are the major contacting region (Fig. S2A). For the R89E mutant, most of the resonances were preserved including those from the α4-α5 residues (Fig. S2B). We thus implemented the same lipophilic PRE approach and observed a slower PRE-induced NMR spectral peak intensity decay for the R89E mutant in bicelles (Fig. S6C), indicating that the replacement of R89 with a glutamic acid weakened the core dimer interaction with the lipid bilayer. Presumably reduced membrane binding is due to the introduction of repulsive negative charges close to negative charges in lipid headgroups.

**Figure 4.**
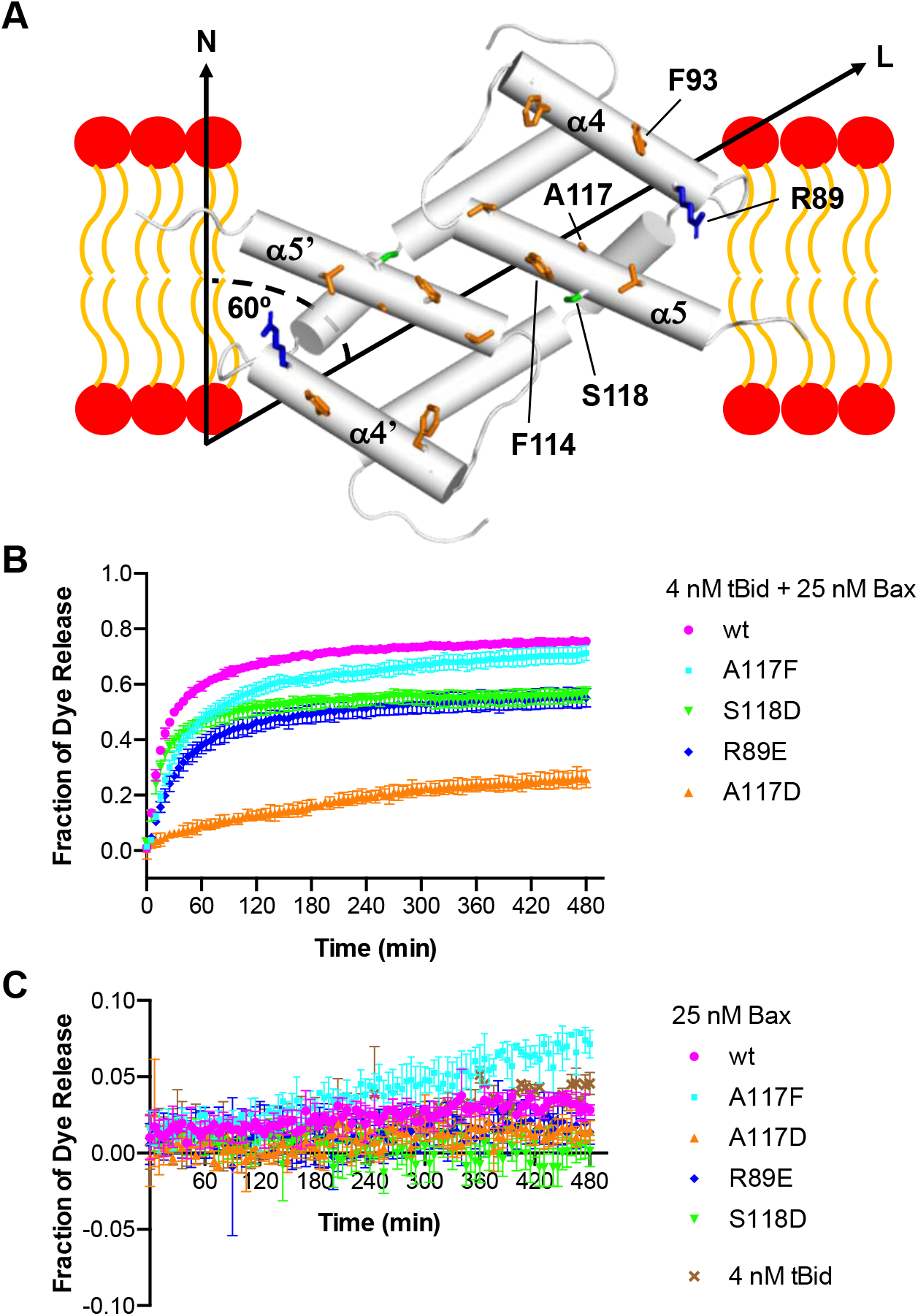
Pore-forming activity of Bax proteins with mutations that would reduce the α2-α5 dimer interaction with membranes. (A) NMR structure of Bax (α2-α5) dimer and lipid bilayer-interacting residues. α helices are shown as cylinders. In the α4 and α5 helices, some of the lipid bilayer-interacting nonpolar, polar and positively charged residues are shown as orange, green and blue sticks, respectively, including the R89, F93, F114, A117 and S118 that were mutated to negatively charged residues to disrupt the interaction with lipid headgroups (red ovals) or acyl chains (orange curves). The longest axis (L) of the dimer is tilted 60° from the bilayer normal (N) to maximize the nonpolar interaction with the lipid acyl chains while allowing R89 to interact with the polar lipid headgroups. (B) Fluorescent dye release from the mitochondria-mimic liposomes by the indicated wild-type (wt) or mutant Bax in the presence (B) or absence (C) of tBid or by tBid alone (C) was measured by quenching the fluorescence of the released dyes by the dye-specific antibodies outside the liposomes during a time course. The fraction of dye release was normalized to that by detergent and shown as means (as symbols) ± standard deviations (SD; as error bars) from three independent experiments.

We further determined the pore-forming activity of the resulting full-length mutant Bax proteins in liposomes that have mitochondrial lipids in the membrane and fluorescent dye conjugated dextran molecules of 10 kDa in the lumen to mimic the mitochondrial protein cytochrome C. The release of the fluorescent molecules from the liposomes was monitored by quenching of the fluorescence by the anti-fluorescent dye antibodies located outside of the liposomes. As shown in Figure 4B, in the presence of 4 nM of tBid protein, 25 nM of wild-type Bax protein released ~75% of the dyes by the end of a 480-min time course. In comparison, the R89E or S118D mutant released ~55% of the dyes, while the A117D mutant released ~25% of the dyes. In contrast, the A117F mutant, as a control for increasing the size of the side chain but not introducing a negative charge, released a slightly lower amount (~70%) of dye than the wild type protein with a slightly slower kinetics. In the absence of tBid, all the proteins released less than 10% of the dyes, demonstrating as expected that this low nM concentration of Bax requires activation by tBid to form pores in the membranes (Fig. 4C). These results clearly show that the mutations which are predicted by our structure-based model to reduce the core dimer interaction with the lipid bilayer reduced the pore formation by the Bax protein.

Since the hydrophobic interaction of the core dimer with the lipid bilayer is extensive and mediated by many residues, mutations of individual residues were expected to have small effects and thereby partially reduce the pore-forming activity. Consistent with this interpretation two additional mutations in the nonpolar α4-α5 surface, F93E and F114E, partially inhibited membrane permeabilization (Fig. S7A). Furthermore, the effects of the R89E, S118D and A117D mutations, detected at 25 nM Bax were greatly reduced by increasing the concentration of Bax. Indeed, the significant difference between the wild-type and mutant proteins in dye release was not detected when 200 nM Bax were used with 4 nM tBid (Fig. S7B). At this high nM concentration, in the absence of tBid, the wild-type protein displayed some auto-activity as it released ~25% of the dyes by the end of the time course (Fig. S7C). Compared to the wild type protein, the S118D and R89E mutants displayed lower auto-activity, whereas the A117D and A117F displayed higher auto-activity (Fig. S7C). Note that the auto-activity of the A117F mutant is higher than the A117D mutant, consistent with the membrane interaction mediated by the core dimer being important for the pore formation by Bax in the absence of tBid.

Bax activation by BH3 proteins is thought to involve unfolding of the soluble structure. Since the A117 and S118 residues are located in the hydrophobic core of soluble Bax protein structure, changing them to bulky and/or charged residues may unfold the structure thereby activating the protein. To test this possibility, we measured the intrinsic fluorescence from the six Trp residues in the Bax protein that are sensitive to their local environment which in turn indicates the folding state of the protein (Fig. S8A). As shown in Figure S8B, the A117D or A117F mutation decreased the Trp fluorescence, suggesting that each mutation increases the access of one or more Trp to water, consistent with the mutation unfolding the protein structure. This unfolding provides an explanation for the high autoactivity of these mutants in the pore-forming assay. In contrast, the S118D mutation increased the Trp fluorescence, suggesting that this mutation does not unfold the protein but changes the conformation thereby reducing the Trp exposure to water (Fig. S8B). As expected, the R89E mutation on the surface of soluble Bax structure did not alter the Trp fluorescence, indicating that this mutation does not change the access of water within the protein structure (Fig. S8B). Consistent with the impact of the S118D or R89E mutation on the protein conformation, these mutations do not increase auto-activation of the protein.

The NMR structure of Bax (α2-α5) dimer bound to the lipid bicelle is different from the crystal structure, suggesting that the core dimer structure is altered by the interaction with the lipid bilayer. To determine the relationship between the core dimer structure and its interaction with membranes, we monitored the dimerization of full-length Bax proteins via the core region in the isolated mitochondria by disulfide crosslinking of Bax mutants with cysteine residues positioned in the core dimer interface (Fig. S9A) (Zhang et al., 2016). As shown in Figure S9B and S9C, the R89E, A117D or S118D mutation reduced the crosslinking of Bax proteins via the L59C in the α2 helix of one protomer and the M79C’ in the α3 helix of the other protomer, and via the two E69C, one in the α2 helix of each protomer. In contrast, the control mutation A117F reduced the Bax crosslinking to a less extent. These data suggest that the mutations that alter the core dimer-membrane interaction also alter the core dimer configuration such that the L59C and M79C or the two E69C are separated by too large a distance to be linked by a disulfide. Because these mutations are not in the core dimer interface (Fig. S9A), they are not expected to reduce the dimerization directly. Note that the mitochondrial association of Bax was largely unaffected by the mutations that reduce the core dimer interaction with the mitochondria (Fig. S9B-C), which is expected since the interactions of other Bax regions with the mitochondria would be retained, in particular, the insertion of the α9 helix into the MOM. Thus, the reduction of Bax crosslinking cannot be attributed to the reduction of Bax concentration in the mitochondria. Therefore, the alteration of the dimer configuration detected by the crosslinking is most likely due to the alteration of the dimer interaction with the membrane.

## Discussion

Since the crystal structure of the Bax core dimer was determined in 2013, different models for its function in Bax pore formation have been proposed (Bleicken et al., 2018; Bleicken et al., 2014; Cosentino and Garcia-Saez, 2017; Czabotar et al., 2013; Uren et al., 2017). In this study, we obtained multiple lines of evidence strongly suggesting that the core dimer forms part of the wall around the pore in the membrane as shown in Figure 5.

**Figure 5.**
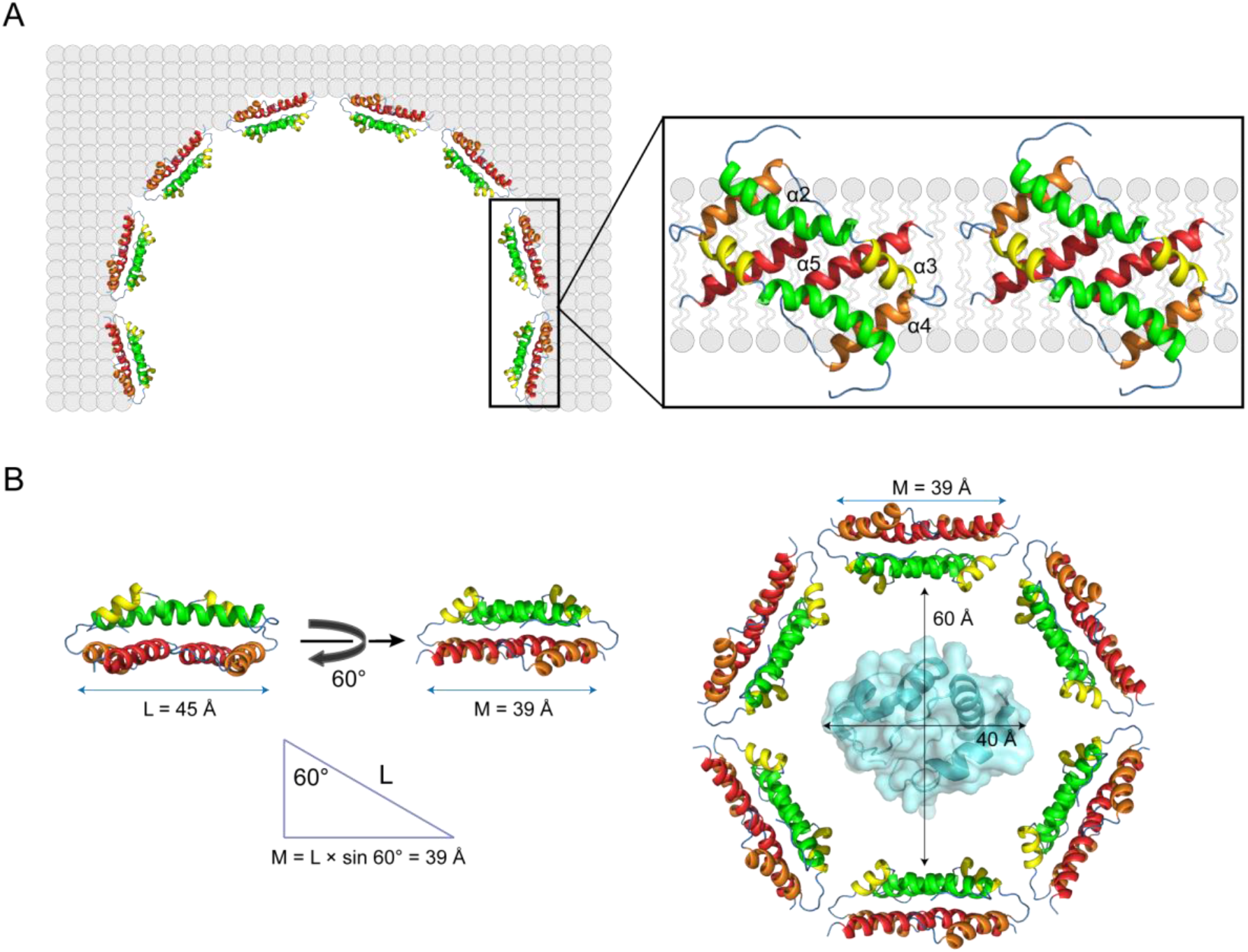
A structural model for an oligomeric Bax pore in the mitochondrial outer membrane. (A) As demonstrated by our study and shown in the model, Bax core dimers form parts of the proteolipid wall that separates the nonpolar membrane from the aqueous pore. Other regions of the Bax oligomer and radially arranged lipids may form other parts of the wall. The diameter of the pore will be determined by the stoichiometry of the Bax oligomer and the number of the radially arranged lipids. (B) In order to release a hydrated cytochrome c from the mitochondria, the diameter of the pore must be > 60 Å, the diameter of cytochrome c plus a water shell. The length of a core dimer along its longest axis (L) = 45 Å. Since the axis tilts 60° from the bilayer normal, a core dimer can cover 39 Å of the lipid bilayer around the pore (M). Assume the pore diameter = 60 Å, and the pore wall only consists of Bax core dimers, six Bax core dimers or 12 Bax proteins are required to form a hexagonal pore that can release cytochrome c. The structure of cytochrome C in the pore was generated from PDB file 3ZCF using PyMOL program.

First, we determined the first structure of the core dimer interacting with a model lipid bilayer using NMR. This structure demonstrates that the core dimer can cover a side of the lipid bilayer using the nonpolar surface formed by the α4-α5 helices that is flattened compared to the crystal structure. This flattening increases the contacting area with the lipid bilayer (Fig. 5A, magnified region). In addition, the longest dimer axis is tilted ~60° from the bilayer normal maximizing the interaction with the lipids, including not only the hydrophobic interactions of multiple nonpolar residues and two polar serines (one from each protomer) with nonpolar lipid acyl chains, but also the ionic or polar interactions of two positively charged arginines (one from each protomer) with the negatively charged or polar lipid headgroups. Since the two arginines are located on the boundary of the nonpolar surface, their interaction with the lipid head groups may dictate the relative orientation of the core dimer to the bilayer normal. While the hydrophobic match between the core dimer structure and the 46-Å thick lipid bilayer formed by DMPC is maximized at the 60° tilt angle, this dimer could match with other bilayers of more or less thickness by a decrease or increase in the tilt angle, respectively. This may explain how Bax permeabilizes multiple membranes of different lipid compositions in vitro (Shamas-Din et al., 2015), and may also permeabilize endoplasmic reticulum, lysosome and other organelles in addition to mitochondria in cells (Bove et al., 2014; Kagedal et al., 2005; Kanekura et al., 2015).

Second, the high-resolution structure of the lipid bilayer-bound core dimer structure enabled us to perform site-specific mutagenesis experiments to examine the contributions of different interactions to the overall pore-forming activity of intact Bax protein in a model mitochondrial membrane that releases a molecule with a size similar to cytochrome C. The results clearly show that not only the hydrophobic interactions of two nonpolar alanines and two polar serines near the center of the nonpolar surface with the lipid acyl chains but the ionic and polar interactions of the two positively charged arginines on the boundary with the lipid headgroups are important driving forces for the Bax pore formation. The functional impact of these site-specific mutations on the intact Bax protein further supports the pore wall-forming function that we proposed based on the NMR structure of the bilayer-bound core dimer. The involvement of the core dimer in assembly of the pore wall as proposed here is consistent with that proposed by the clamp model based on the low-resolution distance distribution data from EPR (Fig. 1A) (Bleicken et al., 2014). However, the EPR data did not reveal the tilt angle or identify which residues interact with the lipid acyl chains versus the lipid headgroups. Furthermore, our structural-guided mutagenesis study provides the first functional validation of our refined model that places the amphipathic core dimer into the wall that separates the hydrophobic membrane from the aqueous pore (Fig. 5A). Our NMR structure has a flat nonpolar surface that covers the flat edge of the lipid bilayer (Fig. 5A, magnified region). This contrasts with the clamp model in which the α2-α5 dimer has a concave nonpolar surface, proposed to cover the complementary convex surface formed by the micellar lipids at the pore rim (Fig. 1A). Although the micellar lipids were not visible in our NMR analyses, our model suggests that some of them must be displaced by the core dimer, while the remainders localize on both sides of the core dimer covering the rest of membrane edge.

Third, in the in-plane model for the function of the core dimer the nonpolar surface of the dimer contacts the surface of lipid bilayer in order for the bulky side chains of nonpolar residues to wedge through the charged and polar region of the lipid bilayer to reach the hydrophobic core (Fig. 1A). Intercalation of these bulky side chains between the lipids in one leaflet of the bilayer was proposed to asymmetrically expand the leaflet generating sufficient membrane tension to induce a lesion in the membrane that is lined by the radially arranged micellar lipids. While this model satisfies the pore-induction role of Bax protein, there is a thermodynamic barrier to overcome. For the nonpolar bulky amino acid side chains to reach the hydrophobic core of a lipid bilayer, they have to first contact and then penetrate through the charged and polar region formed by the lipid head groups. For this to happen there must be enough free energy to complete this thermodynamically unfavorable process. Moreover, it would result in the charged or polar lipid headgroups being located in the space between the α4 and α5 helices that is mostly filled by interacting nonpolar amino acid side chains. It is not clear how such a structure would be thermodynamically stable. Although our study cannot rule out this in-plane membrane topology of the core dimer, we did not detect such a topology for the core dimer reconstituted into the lipid bicelle that has more than enough area on the top for the core dimer to land “in plane”, indicating that even if it does exist, the in-plane membrane topology is likely a transient state during pore formation, rather than a final stable structure around the pore.

Finally, the configuration of the core dimer appears to be induced upon interaction with the membrane. This conformational change was revealed by comparison of the NMR spectra of the core dimer in the absence and the presence of the lipid bicelles (Fig. 1B), and by comparing the NMR structure of the bicelle-bound core dimer with the crystal structure (Fig. 2B and S4). The conformational change is further supported by our core dimer configuration-specific crosslinking of the Bax proteins, since the mutations that reduce the core dimer interaction with the lipid bilayer also reduce the efficiency of the crosslinking via different points in the dimer interface.

In conclusion, we have determined the first membrane-bound Bax core dimer structure, and validated its function in the pore formation by oligomeric full-length Bax proteins in the mitochondrial membrane. We propose a model (Fig. 5A) that incorporates our structural data and more accurately describes the pore structure as composed of a string of the core dimers as part of the wall encircling the large aqueous conduit that permeabilizes the outer mitochondrial membrane and commits the cell to undergoing apoptosis. We further propose that a minimal of six core dimers formed by 12 Bax proteins are required to form a pore that can release cytochrome C, assuming that the pore is entirely lined by the core dimers (Fig. 5B). The challenge ahead is to determine how Bax dimers are organized along the lining of the pore and the possible distribution of lipids between and or around the dimers. Moreover, structure of the entire pore remain unclear particularly how large diameter pores are formed and maintained and to what extent the pore diameter is variable in mitochondrial outer membranes. Such determinations are critical to finding pharmacologically tractable ways to modulate the pore formation thereby enhancing the death of cancer cells while preserving the life of normal cells (Garner et al., 2019; Gavathiotis et al., 2012; Niu et al., 2017; Reyna et al., 2017).

## Materials and Methods

### Protein expression and purification

*E. coli-codon* optimized DNA encoding residues 53-128 (α2-α5) of human Bax was synthesized (Genewiz), in which two cysteine codons at positions 62 and 126 are replaced by serine codons to improve the stability of the encoded protein. The resulting Bax (α2-α5) DNA was fused with a DNA encoding a Hiss-tag followed by a 3C protease cleavage site (LEVLFQGP), and cloned into a pET28a (+) vector. The resulting plasmid was transformed into *E. coli* BL21 (DE3) cells. The cells were grown in isotopically labeled M9 minimal media at 37 *°C* until optical density at 600 nm reached 0.8-0.9. The fusion proteins were expressed for 18 h at 25 °C after induction with 0.5 mM isopropyl-β-d-thiogalactopyranoside. The cells were collected by centrifugation at 8,000 × g for 30 min at 25 °C, and suspended in *Lysis Buffer* (150 mM NaCl, 1% 3-[(3-cholamidopropyl)dimethylammonio]-1-propanesulfonate (CHAPS), 20 mM Tris-HCl pH 8.0). After the cells were disrupted by sonication, the lysate was centrifuged at 40,000 × *g* for 30 min at 4 °C. The supernatant with the soluble fusion protein was loaded onto a Ni-NTA column, which was sequentially washed with 20 mM imidazole, 40 mM imidazole and 60 mM imidazole in *Lysis Buffer*. The fusion protein was then eluted from the column with 400 mM imidazole in *Lysis Buffer.* The elute was dialyzed against *Dialysis Buffer* (200 mM NaCl, 20 mM MES pH 6.0) at 4 °C for 12 h using 3.5 kDa MWCO Slide-A-Lyzer MINI Dialysis Devices (Thermo Scientific) to remove imidazole and CHAPS. The fusion protein was cleaved by 3C protease in *Dialysis Buffer* containing 1 mM PMSF, 200 mM Na_2_SO_4_ at 4 °C overnight. The resulting tag-free Bax (α2-α5) protein was separated from the His_8_-tag and the protease by Ni-NTA chromatography. The Bax (α2-α5) protein was concentrated and further purified by size exclusion chromatography using a HiLoad^®^ 16/600 Superdex 200 pg column (GE Healthcare) in *Dialysis Buffer.* Based on SDS-PAGE analysis, the peak fractions containing homogenous Bax (α2-α5) protein were pooled and concentrated for NMR measurements. Typical yield of this protein prep was ~12 mg/L of cell culture. The mutant Bax (α2-α5) proteins were similarly expressed and purified.

Plasmids encoding mutant human Bax proteins were generated from the pTYB1 vector-based plasmid encoding the wild type protein (Suzuki et al., 2000) using QuickChange (Thermo Fisher Scientific). These plasmids were used to prepare the recombinant proteins as described (Niu et al., 2017) for liposome membrane permeabilization experiments. The recombinant truncated murine Bid (tBid) protein was also prepared as described (Niu et al., 2017).

### Sample preparation of Bax (α2-α5) in Bicelles

6.78 mg of 1,2-Dimyristoyl-sn-Glycero-3-Phosphocholine (DMPC) (Avanti Polar Lipids) were mixed with 51 μL of 200 mg/mL 1,2-Dihexanoyl-sn-Glycero-3-Phosphocholine (DHPC) (Avanti Polar Lipids) in water to reach a ~1:2 DMPC:DHPC molar ratio. The mixture was subjected to cycles of freeze-thaw, pipetting and vortex until the lipids were fully dissolved and the bicelles were formed homogeneously. The purified Bax (α2-α5) (0.1 mM) was incubated with the bicelles under 20-rpm rotation at 25 °C for 30 min. Five to eight aliquots of 0.1 mM Bax (α2-α5) in bicelles was mixed and concentrated to ~500 μL using Centricon concentrator (EMD Millipore; MWCO, 3.0 kDa). Accordingly, the final sample for NMR measurements contains 0.5-0.8 mM (monomer concentration) Bax (α2-α5), 50-80 mM DMPC, 100-160 mM DHPC, 100 mM NaCl, 10 mM MES pH 6.0, 0.1% NaN_3_, and 10% D_2_O. The DMPC: DHPC ratio was monitored by 1D NMR and the molar ratio (q) was kept 0.5-0.6 for all the NMR samples. DHPC and DMPC with deuterated acyl chains were used to prepare the samples for proteinprotein NOESY experiments.

### SEC-MALS analysis of Bax (α2-α5)

The molecular weight of Bax (α2-α5) protein in solution was determined by size exclusion chromatography-multiple angle light scattering (SEC-MALS) using Agilent 1260 Infinity Isocratic Liquid Chromatography System connected to Wyatt Dawn Heleos II MALS detector, Wyatt Optilab T-rEX Refractive Index detector and ViscoStar III differential viscometer (Wyatt Technology). The chromatography was performed at 25 °C using Superdex 200 10/300 GL column (GE Healthcare) equilibrated with *Dialysis Buffer.* 100 μL of 8.0 mg/mL of Bax (α2-α5) was injected into the column and eluted with *Dialysis Buffer* at a flow rate of 0.5 mL/min. The light absorption at 280 nm, light scattering at 660 nm and refractive index of the elute were simultaneously monitored with the three inline detectors. The data were analyzed by ASTRA software and the molecular weight of Bax (α2-α5) peak fraction was calculated using the three-detector method.

### Characterization of the Bax (α2-α5) oligomeric states by crosslinking

The oligomeric states of the soluble and bicelle-bound Bax (α2-α5) protein were examined by chemical crosslinking using bissulfosuccinimidyl suberate (BS^3^) (Thermo Scientific). The Bax (α2-α5) was first dialyzed overnight against *Crosslinking Buffer* (150 mM NaCl, 100 mM sodium phosphate pH 7.0) at 4 °C. 4 μL of 50 mM BS^3^ solution in water was added to 50 μL of 80 μM soluble or bicelle-bound Bax (α2-α5) protein, respectively. The crosslinking reaction was performed at 25 °C for 30 min and then quenched by 50 mM Tris-HCl pH 8.0.

### NMR data acquisition, processing and analysis

All NMR experiments were conducted on Bruker spectrometers operating at ‘H frequency of 600 or 900 MHz and equipped with cryogenic probes. NMR spectra were processed using NMRpipe (Delaglio et al., 1995) and analyzed using XEASY (Bartels et al., 1995) and CcpNmr (Vranken et al., 2005). Sequence-specific assignment of backbone amide resonance (^1^H^N^, ^15^N, ^13^Cα, and ^13^C’) was accomplished using a series of gradient-selected, TROSY-enhanced triple resonance experiments, including HNCO, HN(CA)CO, HNCA, HN(CO)CA and HNCACB (Salzmann et al., 1998). The NMR data was collected with a uniformly [^15^N, ^13^C, ^2^H]-labeled Bax (α2-α5) protein at 600 MHz. In addition, ^15^N-edited NOESY-TROSY-HSQC experiments (150-ms NOE mixing time) were performed to validate the assignment. Protein aliphatic and aromatic side chain resonances were assigned using a combination of ^15^N-edited NOESY-TROSY-HSQC spectrum and ^13^C-edited NOESY-HSQC spectrum recorded at 900 MHz for a [^15^N, ^13^C]-labeled protein. Stereospecific assignments of methyl groups of leucine and valine were performed for a ^1^H-^13^C HSQC spectrum obtained from a 15% [^13^C]-labeled protein.

To determine the intermolecular distance restraints, a sample containing 1:1 molar ratio of [^15^N, ^2^H] and [^13^C]-labeled Bax (α2-α5) monomer was used to record a ^15^N-edited NOESY-TROSY-HSQC spectrum (250-ms NOE mixing time) to obtain the exclusive NOEs between the ^15^N-attached protons of one monomer and aliphatic protons of the neighboring monomer.

All the NMR experiments were conducted at 25 or 32 *°C* for soluble or bicelle-bound isotope-labeled Bax (α2-α5) sample, respectively, with both samples buffered to pH 6.0. All the lipids were deuterated for the NOESY experiments.

### Structure calculation

Structure was calculated using the program XPLOR-NIH (Schwieters et al., 2018). The crystal structure of Bax (α2-α5) dimer (PDB code: 4BDU) was used to construct an initial model that was then refined against a complete set of NOE restraints (including intramolecular and intermolecular distance restraints) and dihedral restraints using a standard simulated annealing protocol (Bassolino-Klimas et al., 1996). A total of 200 dimer structures were calculated and 15 lowest-energy structures were selected as the structural ensemble.

### Structure validation by PRE measurement

To verify the structures calculated above, PRE analysis was performed to obtain intermolecular distance restraints independent of NOEs. An A82C or S126C mutation was made in Bax (α2-α5) for labeling the protein with MTSL ((1-oxyl-2,2,5,5-tetramethyl pyrroline-3-methyl) methanethiosulfonate). 40 μL of 100 mM MTSL in DMSO was mixed with 1 mL of 0.4 mM [^14^N]-labeled single-Cys Bax (α2-α5) and the labeling reaction was conducted at 4 °C for 12 h. The sample was then dialyzed in *Dialysis Buffer* to remove free MTSL. A sample containing ~1:1 molar ratio of [^15^N] and [^14^N]-labeled Bax (α2-α5) in DMPC/DHPC bicelles (q = 0.5) was prepared as mentioned above in the sample preparation section. The ^1^H-^15^N TROSY-HSQC spectra were recorded before and after the addition of 10 mM sodium ascorbate. The peak intensity ratios of the paramagnetic (I) to the diamagnetic (I_0_) state were calculated for all spectrally dispersed residues.

### Detection of protein-lipid NOE

0.8 mM [^15^N, ^2^H]-labeled Bax (α2-α5) in bicelles containing regular DMPC and deuterated DHPC (q = 0.55) was used to measure protein-lipid NOEs. A ^15^N-edited NOESY-TROSY-HSQC spectrum (200 ms NOE mixing time) was recorded at 32 *°C* and 900 MHz. The NOE spectrum was analyzed using XEASY (Bartels et al., 1995).

### Solvent PRE analysis

The solvent PRE measurements were performed as previously described (Piai et al., 2017). Briefly, a sample of 0.8 mM [^15^N, 85% ^2^H)-labeled Bax (α2-α5) mixed with DMPC/DHPC bicelle (q = 0.6) was titrated with the water-soluble and membrane inaccessible paramagnetic agent Gd-DOTA of 0, 0.5, 2.5, 4.5, 6.5, 8.5, 13.5, 23.5, 28.5, 33.5 and 38.5 mM. A series of ^15^N TROSY-HSQC spectra with 3.5-s recovery delay were recorded at 600 MHz. The spectra were analyzed by CcpNmr (Vranken et al., 2005) to get the ratio of peak intensity in the presence (I) and absence (I_0_) of the paramagnetic agent. The residue-specific PRE_amp_ values were derived by fitting *I/I_0_* vs. [Gd-DOTA] data to an exponential decay (Eq. 1), where τ is decay constant.

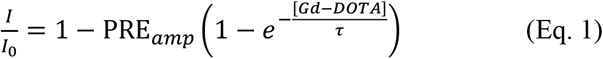

### Lipophilic PRE analysis

The Bax (α2-α5) in bicelle sample was prepared as described above for the solvent PRE experiment. The sample was titrated with the membrane-embedded paramagnetic agent 16-DSA of 0, 0.6, 1.2, 1.8, 2.4, 3.0, 3.6, 4.2 and 4.8 mM. TROSY-HSQC spectra were recorded at each [16-DSA]. The residue-specific PRE_amp_ value was derived by fitting the spectral peak intensity decay as a function of [16-DSA] using a modified (Eq. 1) where [Gd-DOTA] is replaced by [16-DSA]. The standard PPT analysis as described before (Piai et al., 2017) was performed until finding the best fit between the PRE_amp_ values and the sigmoidal function (Eq. 2). Briefly, the projected positions of the amide protons of Bax (α2-α5) protein on the bilayer normal axis were obtained, and the distances of these positions to an arbitrary point (O) on the axis (r_z_) were determined (Fig. S5C). The PRE_amp_ vs. r_z_ data was plotted and fit by (Eq. 2) to determine the position of the Bax (α2-α5) protein relative to the lipid bilayer, in particular, the angle between the longest axis (L) of the Bax (α2-α5) dimer to the bilayer normal axis (N) (Fig. S5D). The lipophilic PRE measurement for Bax (α2-α5) R89E was performed similarly.

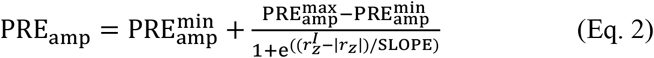

where 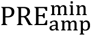 and 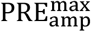 are the minimal and maximal PRE_amp_ possible for a particular protein system, 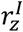 is the distance from the bilayer center to the inflection point at which the PRE_amp_ is halfway between 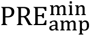 and 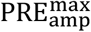), and SLOPE is a parameter for the steepness of the PRE_amp_ curve at the inflection point.

### Liposome permeabilization by Bax

Liposomes were prepared with chicken egg phosphatidylcholine and trans phosphatidylethanolamine, porcine brain phosphatidylserine, soy phosphatidylinositol, and bovine heart cardiolipin (Avanti Polar Lipids) of 47:28:9:9:7 mol%, a lipid composition found in Xenopus oocyte mitochondria (Kuwana et al., 2002), using an extrusion method as described (Tan et al., 2006). Fluorescent dye Cascade blue (CB)-labeled dextrans of 10 kDa (ThermoFisher Scientific) were encapsulated in the liposomes. To monitor the liposomal membrane permeabilization by Bax proteins, the liposomes with 50 μM total lipids were mixed with 12 μg/mL anti-CB antibody (ThermoFisher Scientific) in 250 μL of buffer A (50 mM Na_2_HPO_4_ adjusted to pH 7.4 using ~3 mM citric acid) in a 4 × 4-mm quartz microcell. The initial CB fluorescence intensity F_0_ was measured at 37 °C using ISS PC1 photon counting spectrofluorometer with excitation and emission wavelength set at 400 and 430 nm, respectively. The slit widths of excitation and emission monochromators were 2.4 and 2.0 mm, corresponding to the spectral resolutions of 19.2 and 16 nm, respectively. 25 or 200 nM Bax protein and/or 4 nM tBid protein were added to the microcell. After 3 min of mixing and equilibrating the sample to 37 °C, the CB fluorescence intensity F(t) was measured for 3 sec every 5 min for 480 min to monitor the quenching of the released CB-dextrans by the anti-CB antibodies located outside of the liposomes. At the end, 0.1% Triton X-100 was added to lyse all the liposomes, release all the CB-dextrans and quench all the fluorescence. The final intensity F_T_ was then measured at 37 °C. The fraction of CB-dextran release caused by the proteins equals to the extent of fluorescence quenching by the anti-CB antibody in the presence of the proteins normalized to that in the presence of Triton, i.e., [F_0_-F(t)]/[F_0_-F_T_].

### Tryptophan fluorescence emission spectrum of Bax proteins

The tryptophan fluorescence spectrum of 1 μM wild-type or mutant Bax protein in buffer A was measured in a 4 × 4-mm quartz microcell at 37 °C with the excitation and emission wavelengths set at 295 nm and from 315 to 385 nm, respectively, and the same slit widths of excitation and emission monochromators described above. The fluorescence intensity at each nm of emission wavelength was measured for 1 s. Three technical repeats of the spectral measurement were performed for each sample.

### Disulfide crosslinking of Bax in mitochondria

[^35^S]Met-labeled Bax L59C/M79C or E69C protein without or with the additional mutation as indicated in Figure S9 was synthesized in vitro using TNT coupled wheat germ extract system (Promega). As previously described (Zhang et al., 2016), the resulting proteins were targeted to the isolated Bax^-/-^/Bak^-/-^ mitochondria by Bax BH3 peptide. The mitochondria-bound proteins were separated from the soluble proteins by centrifugation and oxidized with copper (II) (1,10-phenanthroline)3 (CuPhe) in induce disulfide crosslinking. The crosslinked Bax dimer was separated from the monomer by non-reducing SDS-PAGE and detected by phosphor-imaging using Fuji FLA-9000 multi-purpose image scanner. Intensities of both dimer and monomer bands in the phosphor-image were measured using Fuji Image Analysis Software Multi Gauge. The dimer to monomer ratio was calculated from the intensities. The mitochondrial association efficiency was also calculated from intensities of Bax proteins in the mitochondrial and soluble fractions.

## Acknowledgements

We thank the staffs from Large-scale Protein Preparation, Nuclear Magnetic Resonance and Mass Spectrometry Systems at National Facility for Protein Science in Shanghai, Zhangjiang Laboratory, China for technical support and assistance in data collection and analysis. This work was supported by grants from National Key R&D Program of China (2017YFA0504804), Key Research Program of Frontier Sciences, CAS (QYZDB-SSW-SMC043) to B. O., by grants from US National Institutes of Health (R01GM062964), OCAST (HR16-026) and Presbyterian Health Foundation to J.L., by an Institutional Development Award from the National Institute of General Medical Sciences of US National Institutes of Health (P20GM103640), and by a foundation grant from the Canadian Institutes of Health Research (FDN 143312) to D. W. A. D. W. A. holds the Tier 1 Canada Research Chair in Membrane Biogenesis.

## Author contributions

B. O., J. L. and D.W. A. conceived and directed the project. B. O. designed NMR experiments. B. O. and J. L. designed mutations and functional assays. F. L., Y. Y., L. Z. and L. D. prepared Bax (α2-α5) proteins for NMR. F. L., Z. L. and M. W. collected NMR data. F L., M. W. and B. W. analyzed the NMR data and determined the Bax (α2-α5) structure with the help of J. C. and B. O.. F. L. and A. P. analyzed PRE data and built the structural model for bicelle-bound Bax (α2-α5). F. Q. and J. D. R. prepared Bax proteins, performed fluorescence experiments and analyzed the data with J. L. Z. Z. generated Bax mutant plasmids, performed crosslinking experiments and analyzed the data with J. L. B. O., J. L. and D. W. A. interpreted the data and wrote the paper. All authors edited the paper.

---

**Figure S1, Related to Figure 1.**
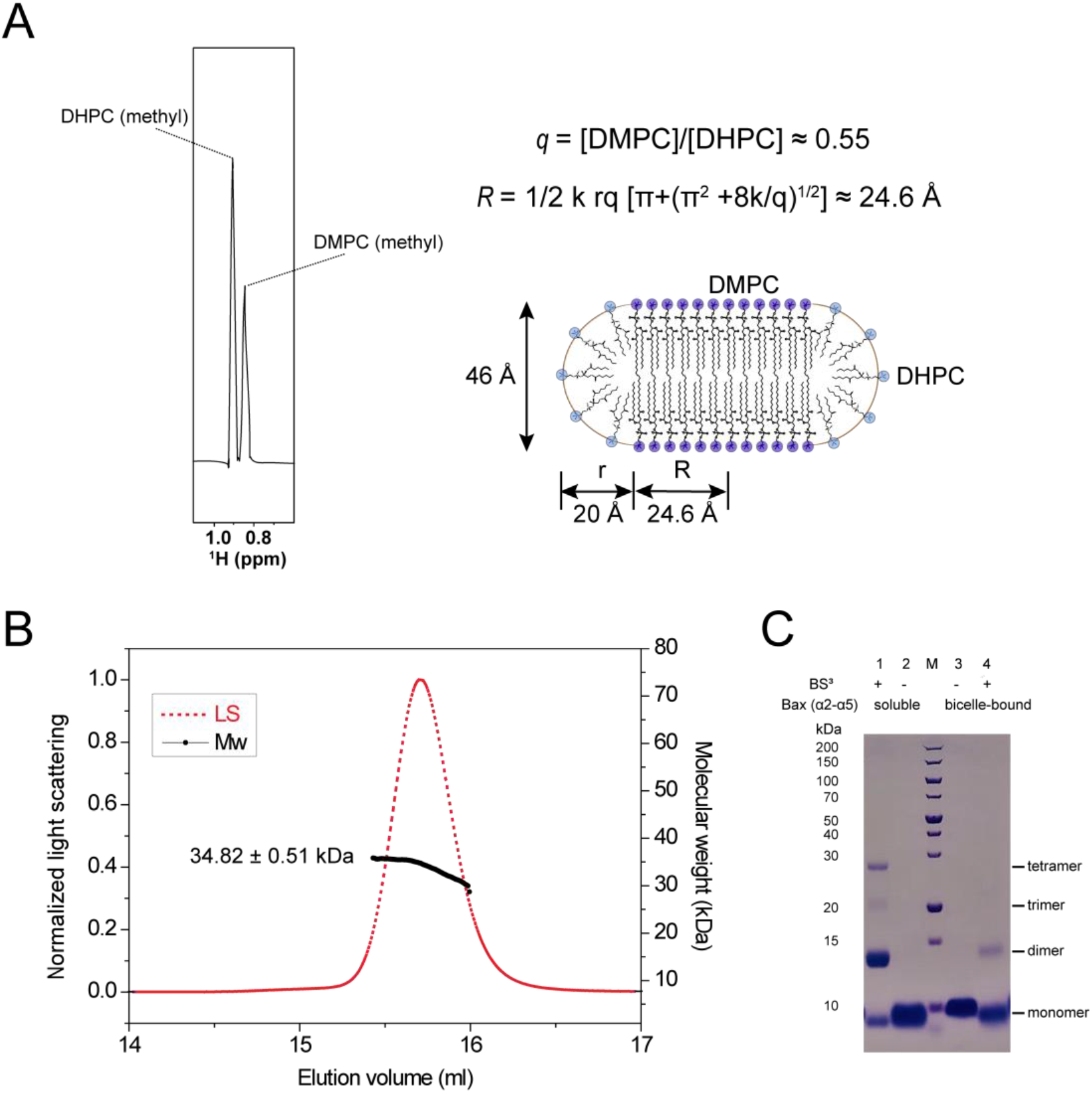
Purification and preparation of Bax (α2-α5) in bicelles. (A) Characterization of the lipid bicelles used in NMR experiments. The molar ratio of DMPC to DHPC (q) in the bicelles determined from the one-dimensional ‘H spectrum on the left is ~ 0.55. The bicelle structure is illustrated with the planar lipid bilayer formed by DMPC and the micellar lipid rim formed by DHPC. The equation describes the radius of the planar region of the bicelle (R) as a function of the molar ratio of DMPC to DHPC (q), where k is the ratio of the head group area of DMPC to that of DHPC, and r is the radius of the rim. In this case, q = 0.55, k = 0.6, and r = 20 Å, thus, R = 24.6 Å. (B) SEC-MALS analysis of the purified Bax (α2-α5) protein. The normalized light scattering (LS) and molecular weight (MW) are shown on the left and right axes, respectively. The black curve represents the calculated MW of the peak fractions of the eluted Bax (α2-α5) with the mean MW and SD indicated. The theoretical MW for Bax (α2-α5) monomer is 9137.43 Da. (C) Chemical crosslinking analysis of oligomerization of the bicelle-bound and soluble Bax (α2-α5) samples used in NMR experiments. The samples from the reactions with or without crosslinker BS^3^ were analyzed by SDS-PAGE and Coomassie Blue staining. The molecular weights for the protein standards in lane M are indicated on the left side. The positions for the monomer and multimers are indicated on the right side.

**Figure S2, Related to Figures 1 and 2.**
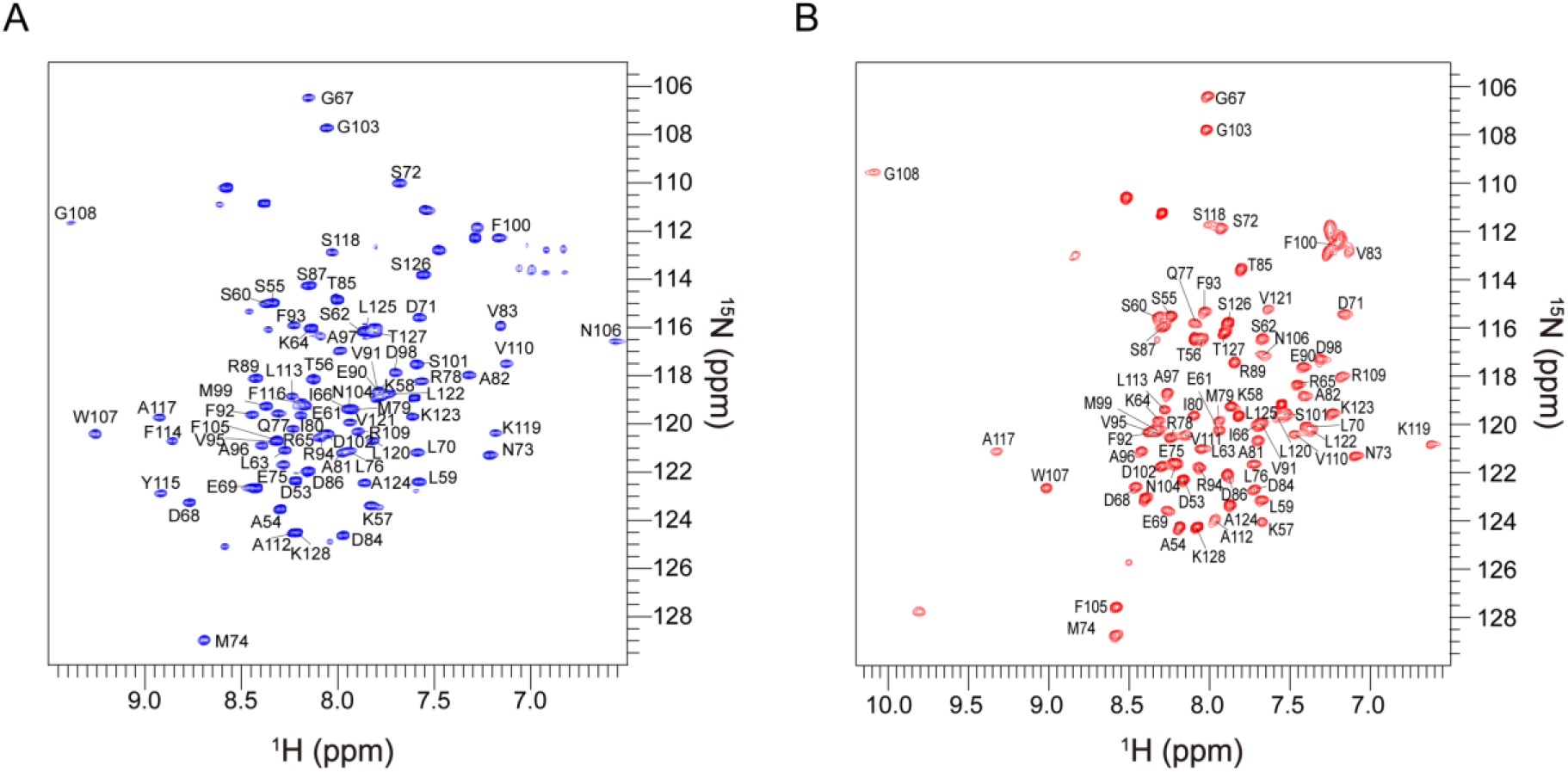
NMR spectra of Bax (α2-α5) in bicelles and solution with assignments. (A) 2D ^1^H^15^N TROSY-HSQC spectrum of the uniformly [^15^N, ^13^C, ^2^H]-labeled Bax (α2-α5) protein in DMPC/DHPC bicelles (q = 0.5) recorded at ^1^H frequency of 600 MHz. The backbone amide resonances were assigned to the indicated residues. (B) 2D ^1^H-^15^N TROSY-HSQC spectrum of the uniformly [^15^N, ^13^C, ^2^H]-labeled soluble Bax (α2-α5) protein recorded at ^1^H frequency of 600 MHz. The backbone amide resonances were assigned to the indicated residues.

**Figure S3, Related to Figure 2.**
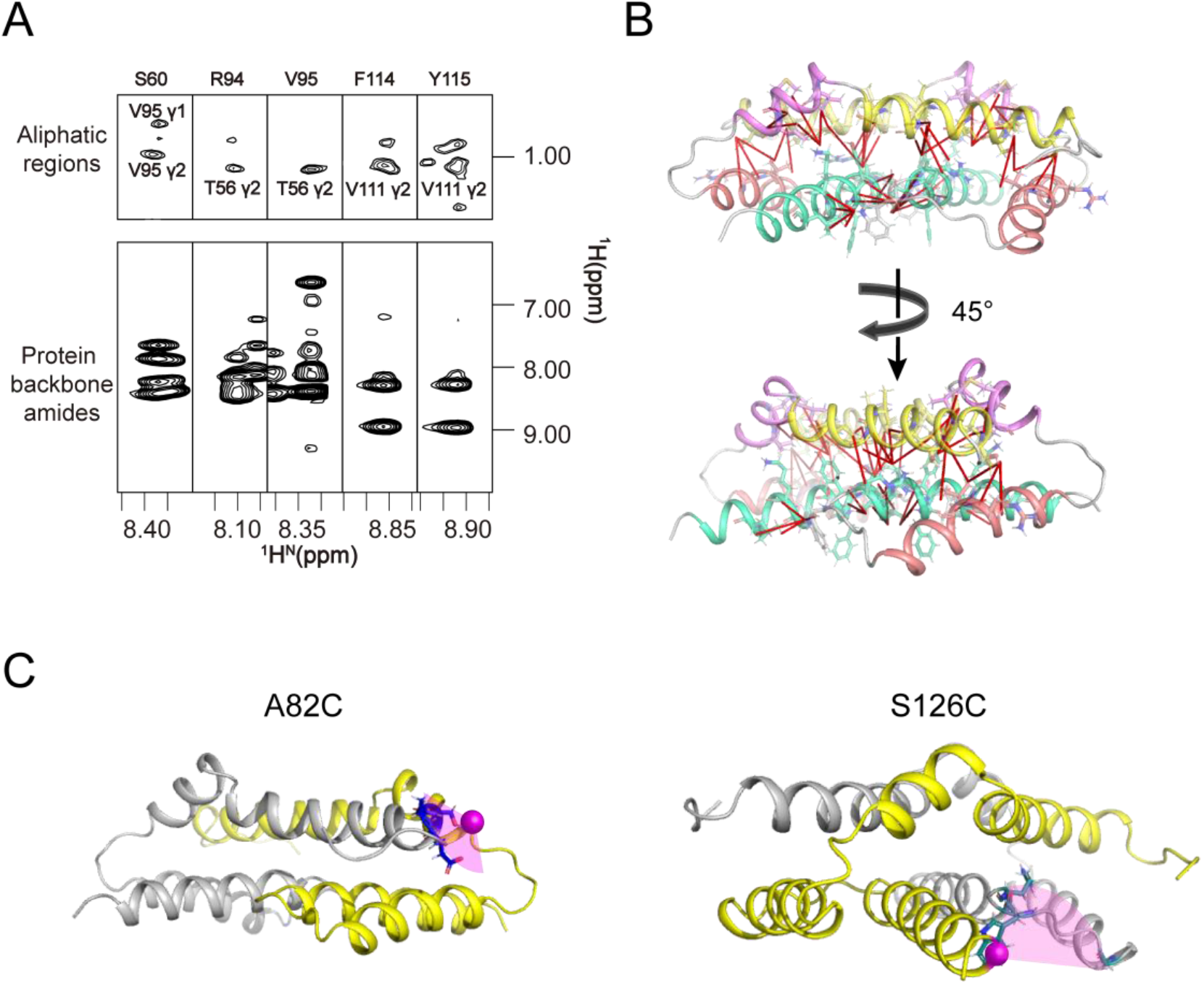
Analysis of intermonomer restraints in Bax (α2-α5) in bicelles. (A) The representative NOE strips taken from 3D ^15^N-edited NOESY-TROSY spectrum recorded at 900 MHz with 250-ms NOE mixing time. The use of mixed isotope-labeled monomers containing 50% [^15^N-, ^2^H]-labeled protein and 50% uniformly [^13^C]-labeled protein permitted the assignment of intermonomer NOEs. The control spectrum from the sample with 100% [^15^N, ^2^H]-labeled protein (data not shown) demonstrated unambiguously that the crosspeaks in the aliphatic regions are the intermonomer NOEs between the backbone amide and the sidechain methyl protons. (B) Sideview of the Bax (α2-α5) dimer structure with all intermonomer NOE-derived distance restraints used in structure determination shown as red lines. (C) Interchain PRE analysis of Bax (α2-α5) in bicelles. Mapping the PREs onto the structure of Bax (α2-α5) in bicelles, showing the backbone amide protons (residue represented by a blue stick in each structure) in chain A (gray) with strong PREs to the nearby spin probe (represented by a magenta sphere in each structure) in chain B (yellow). The sample consisted of an ~1:1 mixture of [^15^N]-labeled Bax (α2-α5) and [^14^N]-labeled Bax (α2-α5) with a MTSL spin probe at A82C (left) or S126C (right).

**Figure S4, Related to Figure 2.**
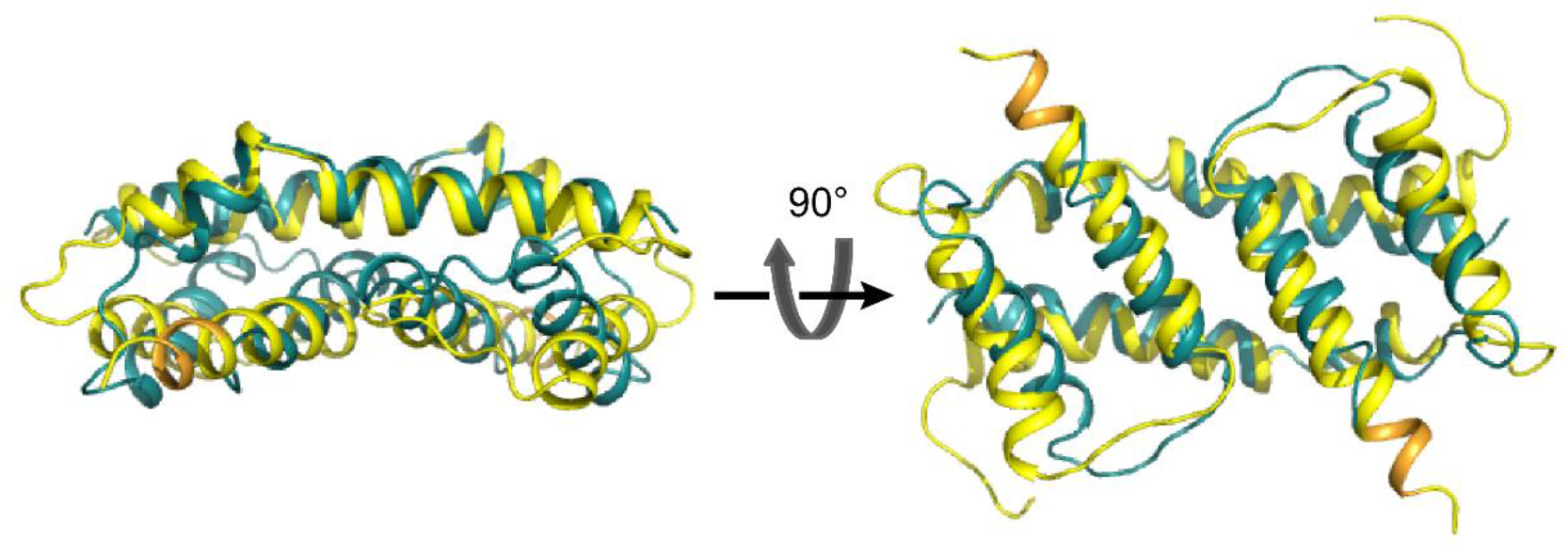
Superposition of Bax (α2-α5) dimer structures in bicelles (yellow) and in crystals (cyan) Three dimensional structure superposition of the structure of Bax (α2-α5) in bicelles determined by NMR (yellow; PDB code: 6L8V) and the structure determnined by crystallography in the absence of membranes (cyan; PDB code: 4BDU).The RMSD value between the superimposed protein backbones is 3.878 Å. Note that the residues from K123 to S126 form an extra α helical turn (orange) in the NMR structure.

**Figure S5, Related to Figure 3.**
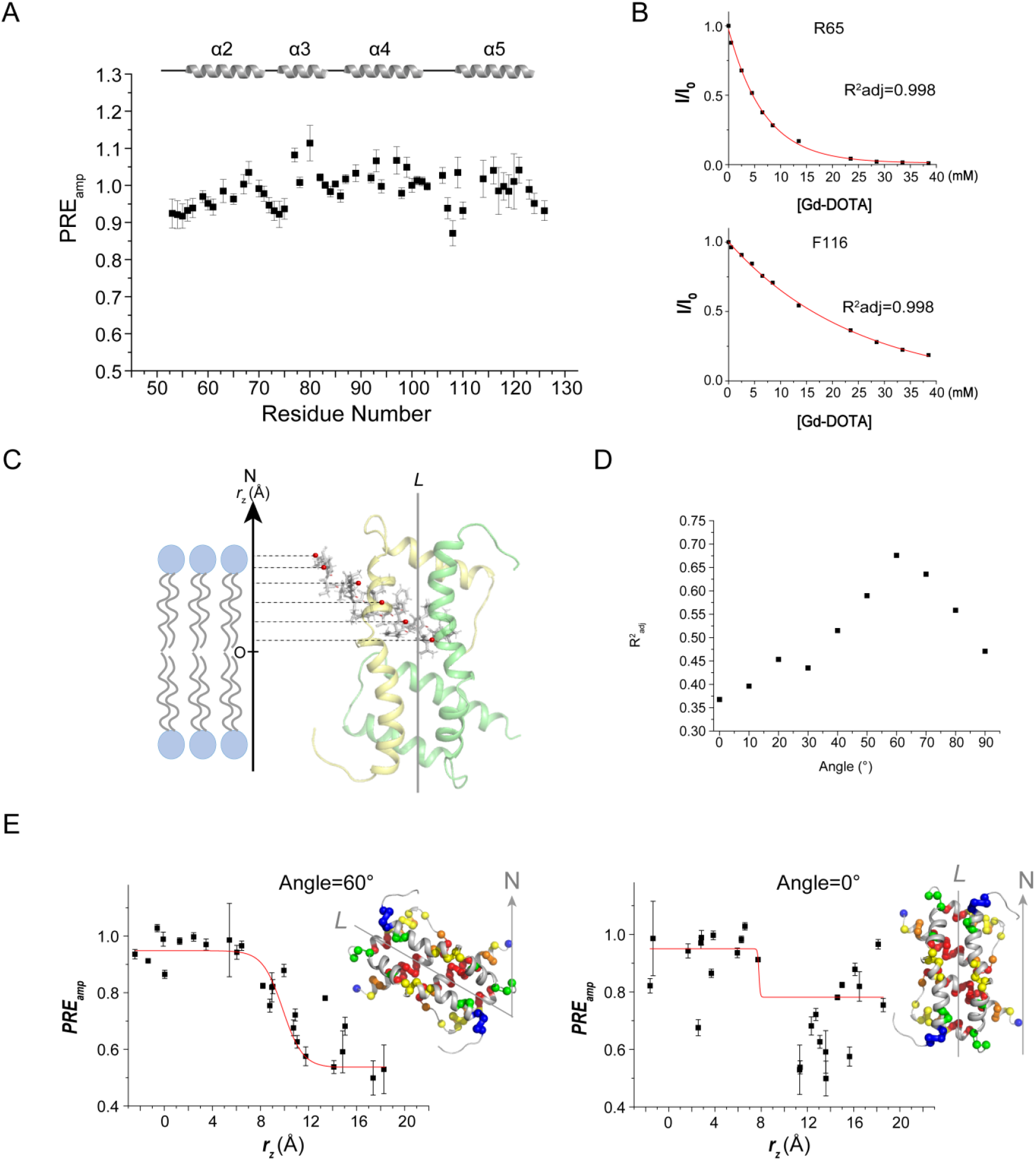
PRE analysis to determine the interaction of the Bax (α2-α5) dimer with the lipid bicelle. (A) Residue-specific PRE_amp_ of Bax (α2-α5) in bicelles derived from Gd-DOTA titration. (B) The ratio of the ^1^H-^15^N TROSY-HSQC spectral peak intensities in the presence (I) and absence (I_0_) of Gd-DOTA for residue R65 in the α2 helix or F116 in the α5 helix versus [Gd-DOTA]. (C) The NMR structure of the Bax (α2-α5) dimer with the longest axis (L) parallel to the lipid bicelle normal axis (N). The amide protons, for which PRE_amp_ has been determined, are shown as red spheres. The dashed lines point to the projected positions of these amide protons on the bilayer normal axis. The distances of these positions to an arbitrary point (O) on the axis are the r_z_ values used in the sigmoidal fitting shown in panel D. (D) R^2^_adj_ versus Angle plot. The quality of the fitting (R^2^_adj_) of the PRE_amp_ (r_z_) data using the symmetric sigmoidal function (Eq. 2) increases as the Bax (α2-α5) dimer rotates closer to the correct orientation relative to the bilayer, which is measured by the Angle between the L and N axes. (E) PRE_amp_ versus r_z_ plots. The structures on the right illustrate two orientations of the Bax (α2-α5) dimer (L) relative to the bilayer normal (N), which result in different r_z_ values for the amide protons, and hence, affect the fitting of the PRE_amp_ vs. r_z_ data using the symmetric sigmoidal function (Eq. 2) as shown by the red fitting curves in the PRE_amp_ (r_z_) plots. The best fitting is achieved at the angle of 60°, whereas the worst fitting is achieved at the angle of 0°.

**Figure S6.**
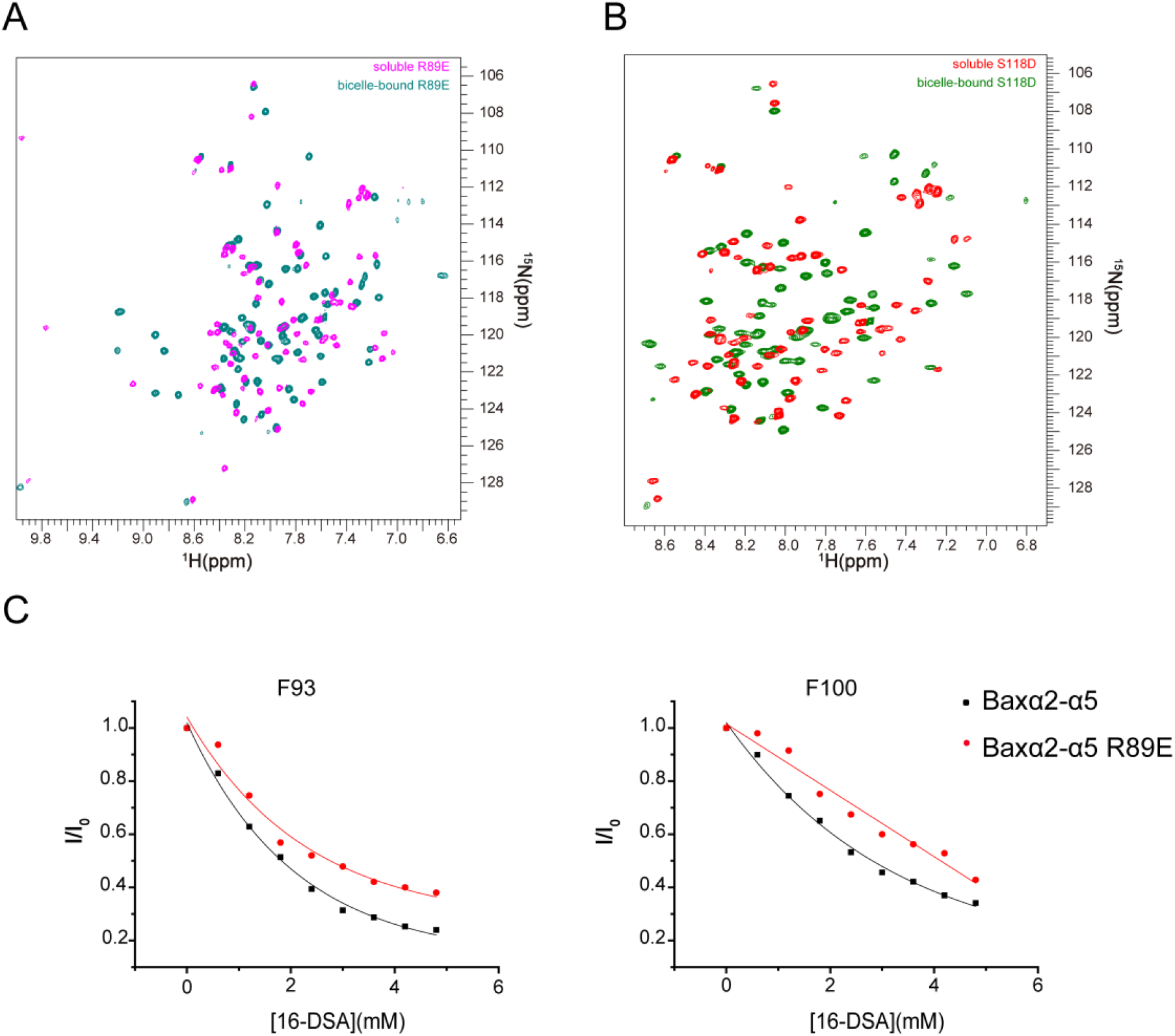
Interactions of Bax (α2-α5) mutants with lipid bicelles. (A-B) Superimposed 2D ^1^H-^15^N TROSY-HSQC spectra of (A) bicelle-bound Bax (α2-α5) S118D (green) and soluble Bax (α2-α5) S118D (red) or (B) bicelle-bound Bax (α2-α5) R89E (teal) and soluble Bax (α2-α5) R89E (magenta) recorded 600 MHz. (C) Spectral peak intensity decay (I/I_0_) for F93 and F100 caused by PRE agent 16-DSA, determined from a series of ^1^H-^15^NTROSY-HSQC spectra of bicelle-bound Bax (α2-α5) or the R89E mutant recorded at different [16-DSA]. I/I_0_ = the peak intensity in the presence 16-DSA/that in the absence of 16-DSA.

**Figure S7, Related to Figure 4.**
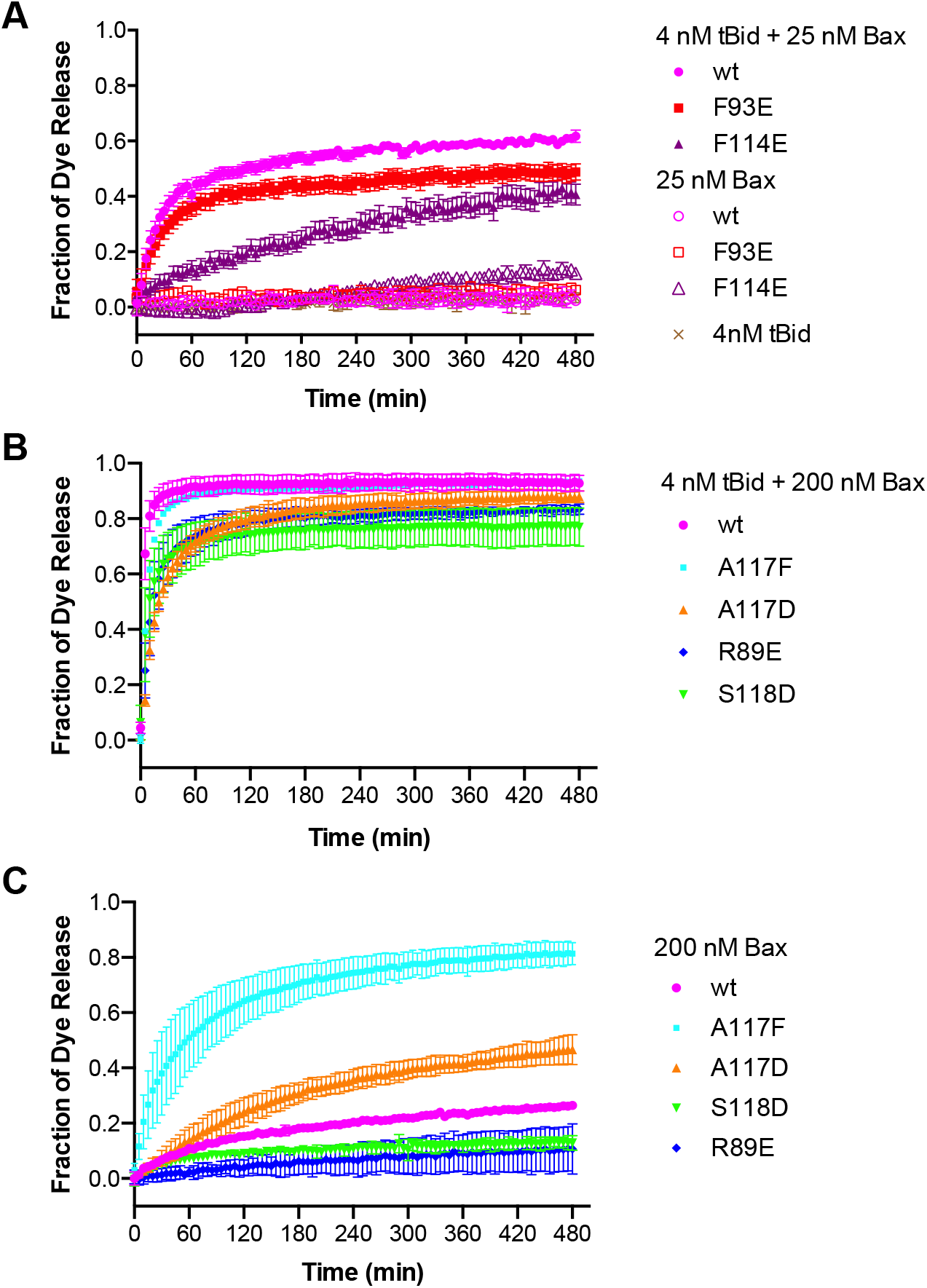
Pore-forming activity of Bax mutants at low or high concentration. (A-C) The fluorescent dye release from mitochondrion-mimic liposomes by 25 or 200 nM wt or mutant Bax in the presence or absence of 4 nM tBid as indicated was measured during a time course by fluorescence quenching. The fraction of dye release was normalized to that by detergent and shown as means (symbols) ± SD (error bars) from three independent experiments.

**Figure S8, Related to Figure 4.**
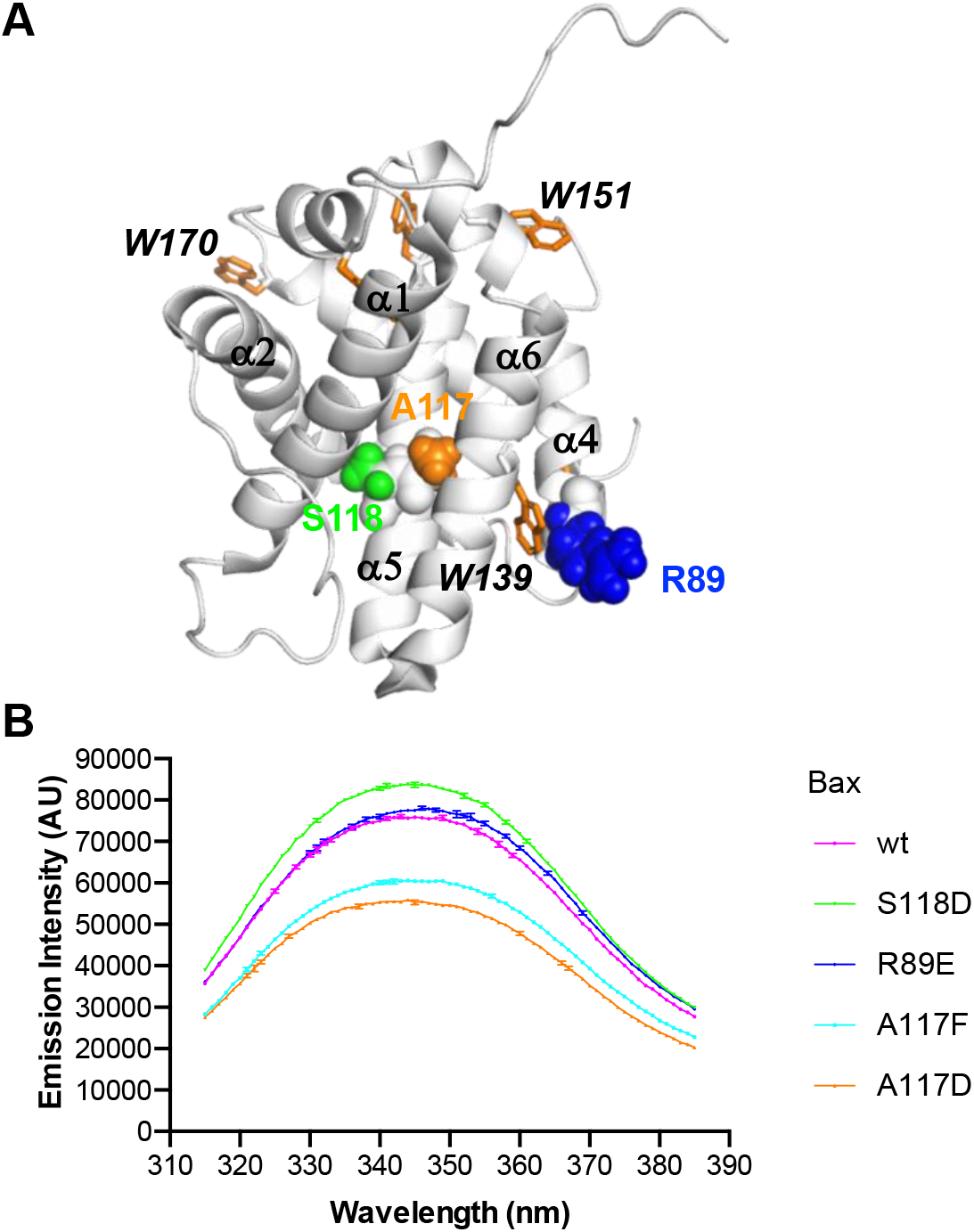
Tryptophan emission spectra of Bax proteins. (A) Bax protein structure. A ribbon diagram of the NMR structure of soluble Bax (PDB code: 1F16) is shown with some of the nine helices labeled. Some of the six Trp residues are visible and shown as orange sticks. R89 on the protein surface, and A117 and S118 in the protein core are shown as blue, orange and green spheres, respectively. (B) Trp emission spectra of the indicated Bax proteins in solution. The data shown are means (lines) ± SD (error bars) from three technical repeats of one experiment.

**Figure S9, Related to Figure 4.**
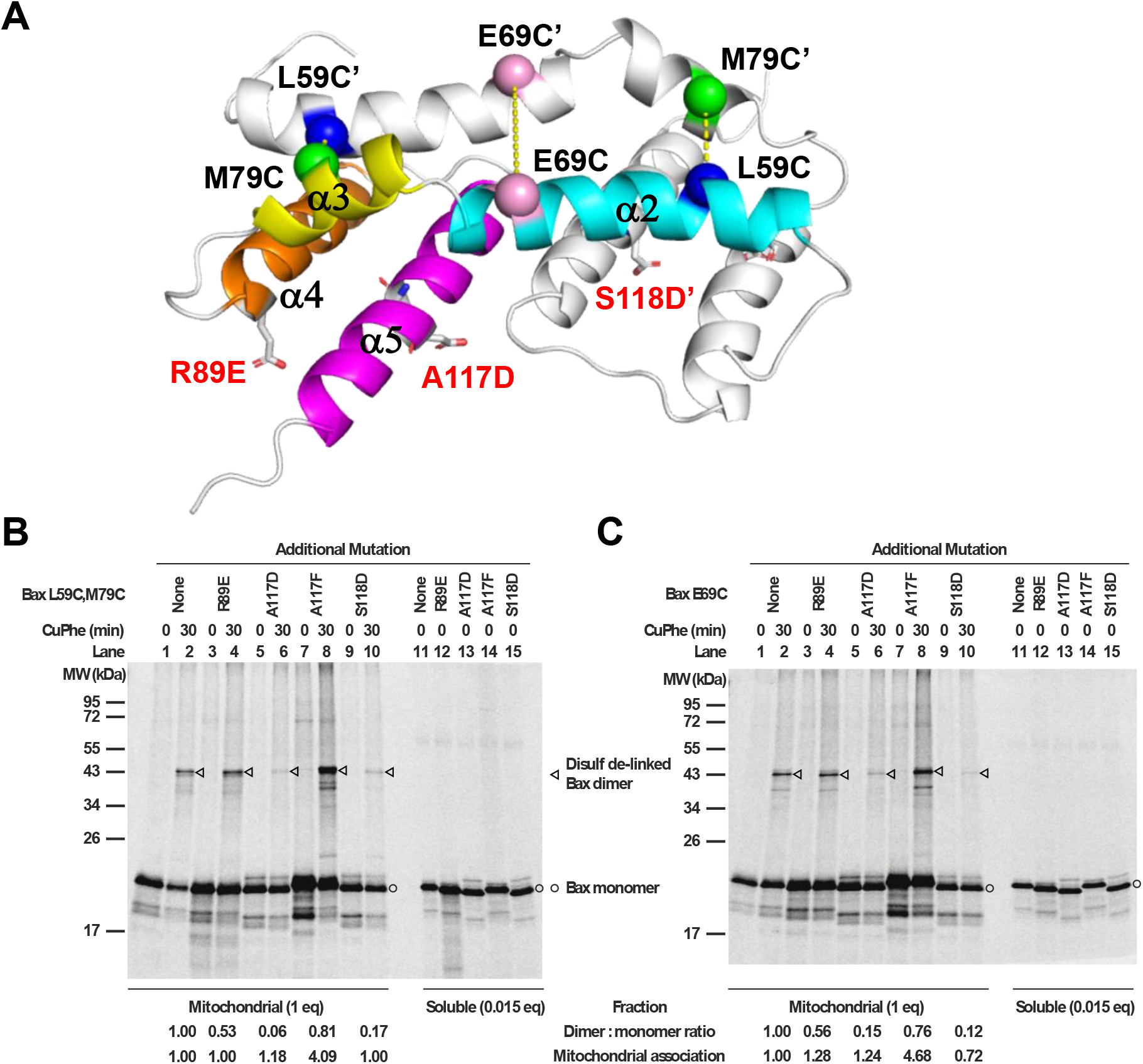
Effect of membrane-binding defective mutations on the Bax core dimerization. (A) A ribbon diagram of the NMR structure of bicelle-bound Bax (α2-α5) dimer. One protomer is colored with the helices labeled. The C_α_ carbons of the residue pairs that were replaced by Cys pairs for disulfide crosslinking are shown as spheres linked by dashed lines. The R89E, A117D and S118D mutations are represented by sticks. (B-C) [^35^S]Met-labeled Bax proteins with two Cys at position 59 and 79 (B) or one Cys at 69 (C) and an additional mutation, if indicated, to disrupt the α2-α5 core dimer interaction with membranes were synthesized, activated by Bax BH3 peptide, and targeted to the mitochondria lacking endogenous Bax and Bak proteins. The mitochondria-bound proteins were oxidized by CuPhe for 30 min to induce disulfide crosslinking of the two protomers via the Cys pair(s) in the core dimer interface as shown in (A). The radioactive crosslinked Bax dimer was then separated from the monomer by non-reducing SDS-PAGE and detected phosphor-imaging. The representative data from two to four independent experiments are shown. The relative dimer:monomer ratio shown at the bottom of the phosphor-images was determined by the intensities of the dimer and monomer bands of the Cys mutant with the additional mutation, normalized to that of the corresponding Cys mutant without the additional mutation. The mitochondrial association efficiency of each Bax mutant (E) was determined by the summed intensity of the monomer and dimer bands in the mitochondrial fraction that was divided and treated with CuPhe for 0 or 30 min (I_M_), the intensity of the monomer band in the corresponding soluble fraction that was divided by 0.015 to adjust the inequivalent loading between the mitochondrial (1 equivalent (eq)) and soluble (0.015 eq) fractions (I_S_), and the following equation, E = I_M_/(I_M_ + I_S_). The relative mitochondrial association efficiency of each Bax Cys mutant with the additional mutation (E_R_) to that of the Bax Cys mutant without the additional mutation was obtained by dividing the E of the former with the E of the latter, and shown at the bottom of the phosphor-images.

